# Eccentricity-Dependent Saccadic Reaction Time: The Roles of Foveal Magnification and Attentional Orienting

**DOI:** 10.1101/2023.08.08.552339

**Authors:** Yufeng Zhang, Pascal Fries

## Abstract

A hallmark of primate vision is the emphasis on foveal processing, accompanied by frequent saccades that bring the fovea to salient parts of the scene, or to newly appearing stimuli. A saccade to a new stimulus is one of the most fundamental sensory-motor transformations. In macaque monkeys, we show that foveal magnification is not only the reason for saccades, but it also affects the dynamics of saccade initiation. In a task where the monkeys made saccades to peripheral target onsets, saccadic reaction time (SRT) increased with target eccentricity. Notably, we effectively eliminated this increase by scaling the target size according to the foveal magnification factor in the superior colliculus. We repeated the comparison between non-scaled and scaled targets, while changing the task to a delayed saccade task. In this task, the target was presented long before the saccade, and the saccade was triggered by foveal fixation offset rather than target onset, such that target onset long before fixation offset was essentially irrelevant for SRT. In this task, we found that SRT increased with target eccentricity, with similar rate for both non-scaled and scaled targets. Furthermore, this increase survived the addition of a salient distracting flash resetting attention to the foveal. The results obtained with the delayed saccades task are consistent with an attentional scan from the fovea to the target, a recently hypothesized general mechanism of attention.

## Introduction

We make several saccades per second, redirecting our gaze to bring objects of interest into our high-resolution fovea. However, the time it takes to initiate a saccade to a novel target, i.e. the saccadic reaction time (SRT) can vary significantly, even when the task is simple and highly repetitive. Part of the variability comes from the size of the impending saccade. The relationship between SRT and the saccade size has a characteristic bowl shape with long SRTs for the shortest (<1 degree of visual angel [dva]) and longest (>10 dva) saccades, and shorter SRTs for medium-sized saccades^1–5^. Within these medium-sized saccades the relationship between SRT and target eccentricity remains unclear and seems task-dependent^1,4,6–9^. Yet, these medium-sized saccades between 2 and 10 dva constitute the most commonly executed saccades in daily life^10,11^ and in laboratory settings, where SRTs are widely used as a tool to characterize the cognitive process of interest^12–14^.

Previous studies describing the relationships between SRT and saccade size often utilized a Step task. In a Step task, the fixation dot steps into the periphery and becomes the saccade target. This design usually comes with a significant confounding factor: saccade targets that are physically identical at various positions vary in visibility with corresponding saccade sizes^15^. This is partially due to the fact that a visual stimulus of a given physical size undergoes foveal magnification early in the visual processing pathways^16–18^ and, therefore, has a decreasing drive for increasing eccentricity or saccade size, respectively^19^. At the single-neuron level, this decreasing drive is correlated with the stimulus occupying a smaller proportion of the receptive field (RF), the size of which increases as the RF moves away from the fovea^16,17,20^. Further, accompanied by foveal magnification and the corresponding change in RF sizes, neurons representing the visual periphery prefer lower spatial frequency components as compared to neurons representing the fovea^21^. Given that a dot is a broad-spectrum stimulus, its spectral components will be amplified differentially at different eccentricities. In addition, low background lighting commonly employed in previous studies might have resulted in the target onset signal being too strong, such that the SRT showed a floor effect, which might have masked the effect of target eccentricity^22^. It is, therefore, important to control the strength of the afferent signal at different eccentricities to draw conclusions about the dependence of SRT on target eccentricity *per se*.

Besides physical stimulation, the nature of the impending saccade, as determined by the specific task structure, can also influence the relationship between SRT and target eccentricity. Recently, Hafed and Goffart^4^ reported a clear effect of increasing SRT with target eccentricity in a visually guided saccade task. Despite the confounding factors mentioned earlier, when the same stimulus set was used for a *delayed* visually guided saccade task, the observed increase of SRT with eccentricity became much less prominent. This is interesting because the main difference between these two tasks is that in the visually guided saccade task, the response saccade was exogenously driven, dominated by the transient onset of the target stimulus; in contrast, in the *delayed* visually guided saccade, the response saccade was endogenously driven, triggered by fixation dot offset, far away from the target stimulus. The observation that the SRT in these two tasks exhibited different patterns of eccentricity dependence motivated us to investigate these two types of saccades (endogenous saccades and exogenous saccades) separately.

Given the same physical conditions and task structure, SRTs across trials can still exhibit high variability^23,24^. The participant can very commonly make a response saccade with SRT as fast as 150 ms in one trial and as slow as 300 ms in another trial. Such high variability requires many trials to reveal a genuine but modest shift of the SRT distribution. For example, more than 200 trials per condition are needed to reveal a shift of 10 ms for a typical SRT distribution (calculated with an ex-Gaussian peaked at 200 ms with a standard deviation of 38 ms; two sided Mann-Whitney U test). Such trial counts per condition were generally not obtained in previous studies.

The current study aims at addressing the aforementioned issues in macaque monkeys, because they offer a rich history of studies of oculomotor control and attention, and also the potential to use the current results to motivate future circuit investigations. We selected two sets of stimuli as saccade targets located between 2 and 10 dva (Figure 1A). One set is Equal, having the same physical size at different eccentricities. The second set is Scaled based on the foveal magnification in the superior colliculus (SC), aiming to equalize the targets’ corresponding afferent input strength to the SC across eccentricities^17^. To avoid a flooring effect due to high input strength, in all tasks, we presented the target on a gray background. We used the same stimuli sets in both, a Step (visually guided saccade) task where the response saccade is exogenously driven by the target onset, and a Delayed (delayed visually guided saccade) task where the response saccade is endogenously triggered by the fixation dot offset (Figure 1B-C). Because the scaling is supposed to mainly affect the target onset transient, we expect to see the scaling to reduce SRT significantly in the Step task but to affect SRT minimally in the Delayed task. Additionally, in the Delayed task, we incorporated an extra condition including a transient attention-capturing distractor flash to investigate whether the eccentricity-dependent SRT changes for the endogenously driven saccades are consistent with an attentional scan from the fovea. Last but not least, for each monkey and each condition, we collected more than 200 trials per condition to gain the necessary statistical power.

**Figure 1.**
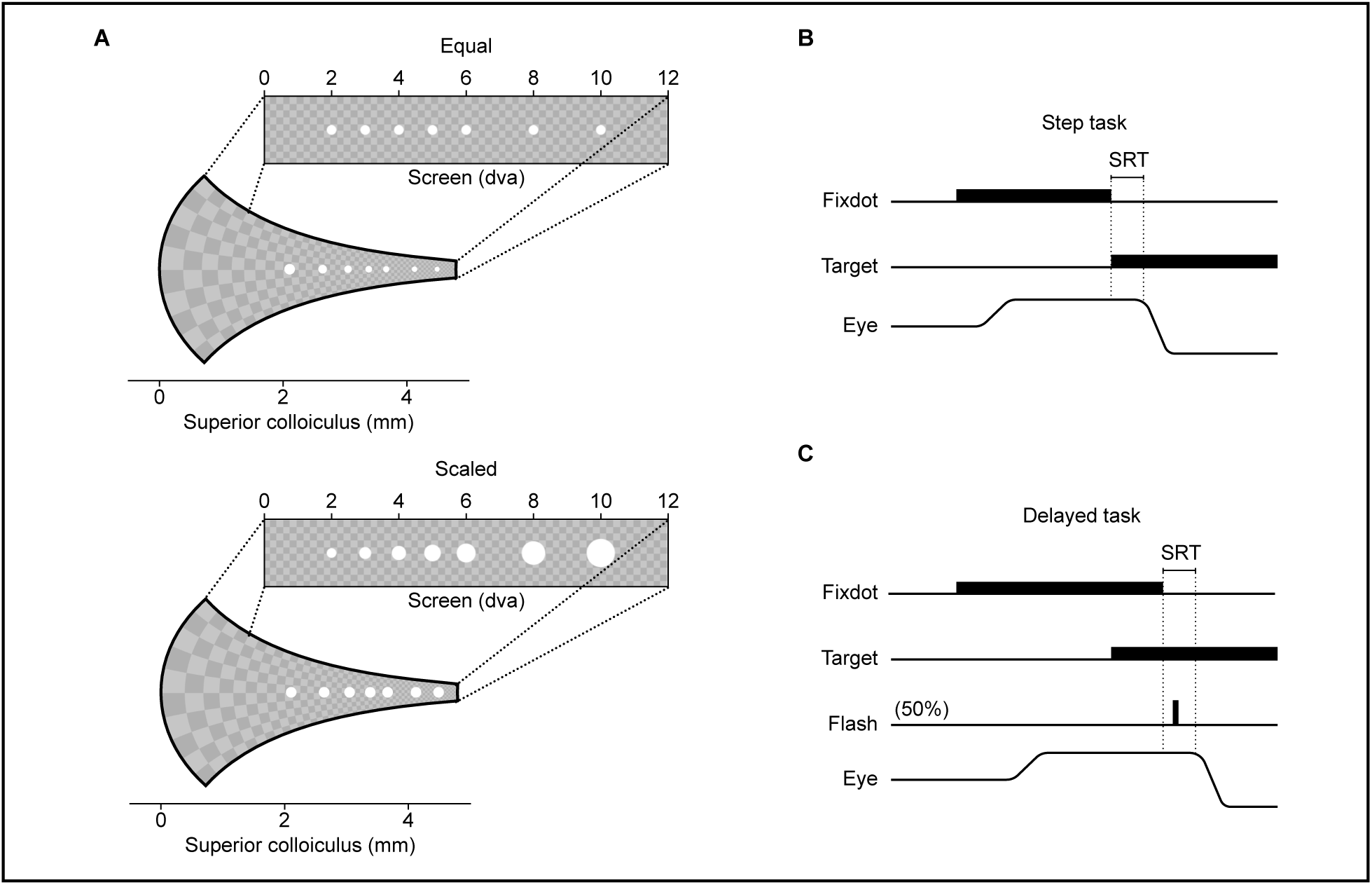
Stimuli and task. (A) Representation of target stimuli on the screen and their corresponding afferent input in the SC map, based on the foveal magnification reported in Chen, et al.^17^. Top: Equal targets at varying eccentricities on the screen exhibit a diminishing size of corresponding afferent input in the SC. Bottom: Scaled targets on the screen have the same size of corresponding afferent input in the SC map. Note that the checkerboard background is shown for illustration purposes and was not presented during the experiment. (B) In the Step task, the fixation dot stepped into the periphery and became the target stimulus. (C) In the Delayed task, the target presentation preceded the fixation dot removal. The animal was required to wait until the removal of the fixation dot before shifting its gaze. In 50% of trials, a transient foveal flash was presented at ≈100 ms after fixation dot removal. The flash had no task relevance and served to capture attention back to the fovea, while the response saccade was planned but not yet executed.

## Results

### Foveal magnification explained eccentricity-dependent SRT increase for exogenously driven saccades

We start with exogenously driven saccades in the Step task (Figure 1B). The hypothesis is that, for exogenously driven saccades, the reported SRT increase with eccentricity^4^ is due to foveal magnification. Specifically, accompanying foveal magnification is the increasing RF size with higher eccentricity in the SC to equalize the size of the active population for the same targets located at different eccentricities^16,17^. Increasing RF size, however, decreases the visibility of the stimulus because it occupies a smaller portion of the (excitatory) RF. This hypothesis also aligns with the recently reported observation that SC neurons at higher eccentricities prefer lower spatial frequencies (corresponding to a larger dot)^21^.

To test this hypothesis, we designed two stimulus sets as saccade targets. The first set, Equal, was similar to what has been used in previous studies^4,6^ (Figure 1A, top). In this set, saccade targets at all positions were white filled circles of diameter 0.1 dva. The second set, Scaled, consisted of filled white circles whose sizes were scaled according to the reported magnification factor in the SC, such that they provided the same afferent input measured in the SC map (Figure 1A, bottom). Both sets consisted of stimuli positioned at 2, 3, 4, 5, 6, 8, and 10 dva to the right of the fixation point along the horizontal median. To facilitate comparison, the target stimulus at 2 dva was of the same size in both sets.

Figure 1B illustrates the task structure of the Step task. In this task, each trial started with the monkey maintaining fixation on a fixation dot (0.1 dva filled white circle) for 800 to 1500 ms. Subsequently, the fixation dot was turned off. At the same video frame, a single saccade target randomly chosen from the combined Equal and Scaled stimuli set was presented. The monkey needed to make a saccade within 500 ms towards the target and hold its gaze on the target for another 800 to 1500 ms. If it completed the trial correctly, a juice reward was provided. Only correct trials were included in the analysis.

The resulting SRT distributions in each condition from one example monkey (HO) are plotted in Figure 2A. The raw data shows that for this monkey: 1) Both location and shape of the SRT distribution varied across conditions. 2) Similar to what has been reported before, the peak of the SRT distributions for Equal targets (blue) gradually increased with increasing target eccentricity. 3) This shift was significantly reduced for Scaled targets (orange). 4) SRTs also became more variable as they became longer. We next quantified these observations.

**Figure 2.**
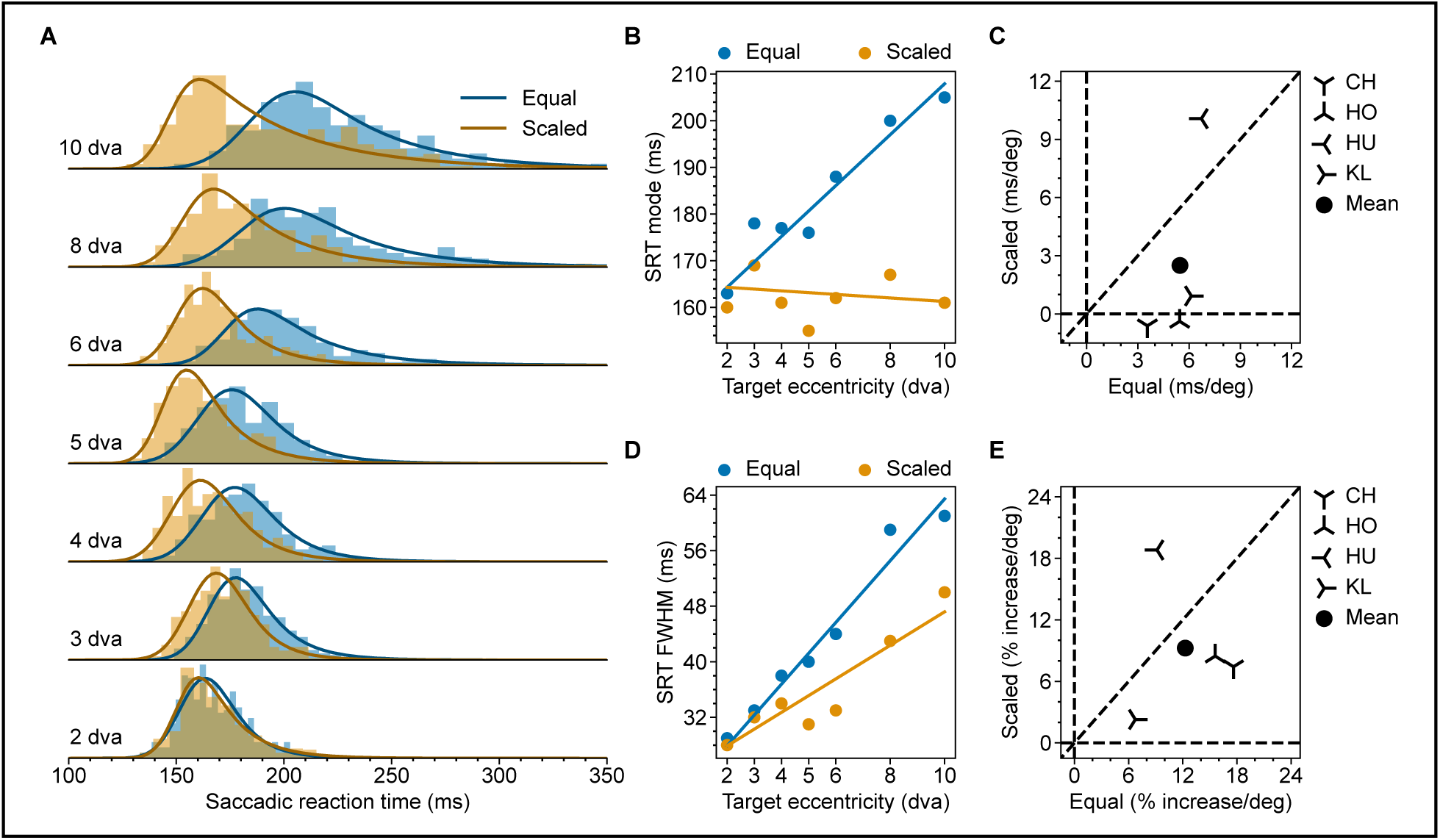
Eccentricity-dependent SRT in the Step task. (A) SRT distributions from monkey HO for Equal (Blue) and Scaled (Orange) targets. Solid lines represent the fitted ex-Gaussian distributions. (B) Multiple linear regression of SRT mode on target eccentricity and scaling for monkey HO. (C) Scatter plot of SRT mode-eccentricity slopes for Scaled vs. Equal targets in all tested monkeys. For each monkey, the slopes are derived from the regression coefficient, as demonstrated in (B). The filled circle shows the mean slopes. (D) Same as (B) but for FWHM. (E) Same as (C) but for FWHM. The raw slopes were converted to percent increase per degree of visual angle. Results for individual monkeys other than HO are provided in the supplementary material (Figure S2-1, S2-2, and S2-3).

To quantify these observations, we first fitted an exponentially modified Gaussian (ex-Gaussian, solid lines in Figure 2A) for each SRT distribution. The ex-Gaussian is the most commonly used parametric function to describe a reaction time distribution^25,26^. We used the mode of each resulting ex-Gaussian fit to represent the typical SRT, as it denotes the most likely occurring value in the distribution. Additionally, we used the full width at half maximum (FWHM) to represent the SRT variability. Having these values, we first examined how the typical SRT varied with saccade size, separately for Equal and Scaled targets. To this end, we fitted a simple linear regression model (*Mode* ∼ β_0_ + β_1_ ∗ (*targetEccentricity* − 2). For monkey HO, SRT mode increased with target eccentricity for the Equal targets but not for the Scaled targets (*Equal*, *CI*_β1,95%_ = [4.33, 5.73], *p* < .001; *Scaled*, *CI*_β1,95%_ = [−0.55, 0.71], *p* = .82). Next, to confirm that scaling reduced the slope of SRT as a function of target eccentricity, we fitted a multiple linear regression model (*Mode*∼ β_0_ + β_1_ ∗ (*targetEccentricity* − 2) + β_2_ ∗ (*targetEccentricity* − 2) ∗ *isScaled*) on the combined data from Equal and Scaled targets (Figure 2B). Scaling indeed reduced the slope (*Combined*, *CI*_β2,95%_ = [−6.45, −5.25], *p* < .001). We repeated the same procedure for the other three monkeys (Figure S2-1, S2-2, and-S2-3). The resulting slopes for each monkey in both conditions are plotted in Figure 2C. Notice that data from three out of four monkeys (CH, HU, and KL) lie very close to the horizontal zero line meaning that, for these monkeys scaling the target stimulus according to the SC foveal magnification factor effectively eliminated the eccentricity-dependent SRT increase. Having only four monkeys, we are essentially limited to the fixed effect analysis and draw inferences on these four monkeys instead of on the population^27^. Fixed effect analysis shows that for these four monkeys, scaling significantly reduced the average slope ( *CI*^β̅̅̅̅^_,95%_ = [−3.42, −2.59], *p* < .001). We repeated the same analysis for the variability of SRT (ex-Gaussian FWHM). Similarly, for monkey HO, FWHM increased with target eccentricity for Equal targets (*Equal CI*_β1,95%_ = [3.43, 5.11], *p* < .001). Scaling also reduced the speed of increase for FWHM but this reduction was not as effective as for the mode (Figure 2D, *Scaled*, *CI*_β1,95%_ = [1.84, 3.24], *p* < .001; *Combined*, *CI*_β2,95%_ = [−2.80, −1.37], *p* < .001). The results for the other three monkeys are presented in the supplementary material (Figure S2-1, S2-2, and S2-3). Fixed effect analysis using averaged percent increase per degree on FWHM agreed with the pattern observed in monkey HO (Figure 2E, *CI*^β̅̅̅̅^_,95%_ = [−4.41, −1.95], *p* = .001).

### Scaling reduced SRT most effectively for low-contrast targets

The results in the previous section suggest that the observed SRT increase with increasing target eccentricity could be explained by foveal magnification. This is presumably because, accompanied by foveal magnification, RF size increases as the RF moves away from the fovea^16,17^. Consequently, a physically identical stimulus closer to the fovea would drive the corresponding neuron stronger in the SC because it occupies a larger proportion of the RF. This stronger stimulus drive would then result in a faster saccade initiation^21^. Also, a stimulus with higher contrast evokes a stronger and faster response throughout the visual hierarchy, leading to a faster reaction time^28–33^. Furthermore, these two bottom-up factors, stimulus size and stimulus contrast, have been shown to interact with each other in determining the magnitude of response transient in the SC^34^. We next investigated whether such interaction between stimulus size and contrast would be reflected in the eccentricity-dependent SRT increase.

To this end, we slightly modified the Step task that we had used so far. Firstly, we reduced the screen background luminance to allow for a wider range of target contrast manipulation. Secondly, at each target location and for each target size, we selected three target luminance levels such that the resulting contrasts were approximately evenly spaced on a logarithmic scale (Weber contrast equals 1.4, 3.1 and 6.9 for Low, Medium and High contrast respectively). Lastly, we limited target locations to 2, 4, and 6 dva to ensure an adequate number of trials for each condition. We collected data from one monkey (HO) for this modified Step task.

We plotted the resulting SRT distributions in Figure 3A-C. At each contrast level, SRT tended to be longer for more eccentric Equal targets. Scaling reduced this increase, and the effect of scaling, in terms of reducing the eccentricity-related SRT increase, appeared to decrease with increasing contrast. As before, we used linear regression to quantify these observations (Figure 3D-K).

**Figure 3.**
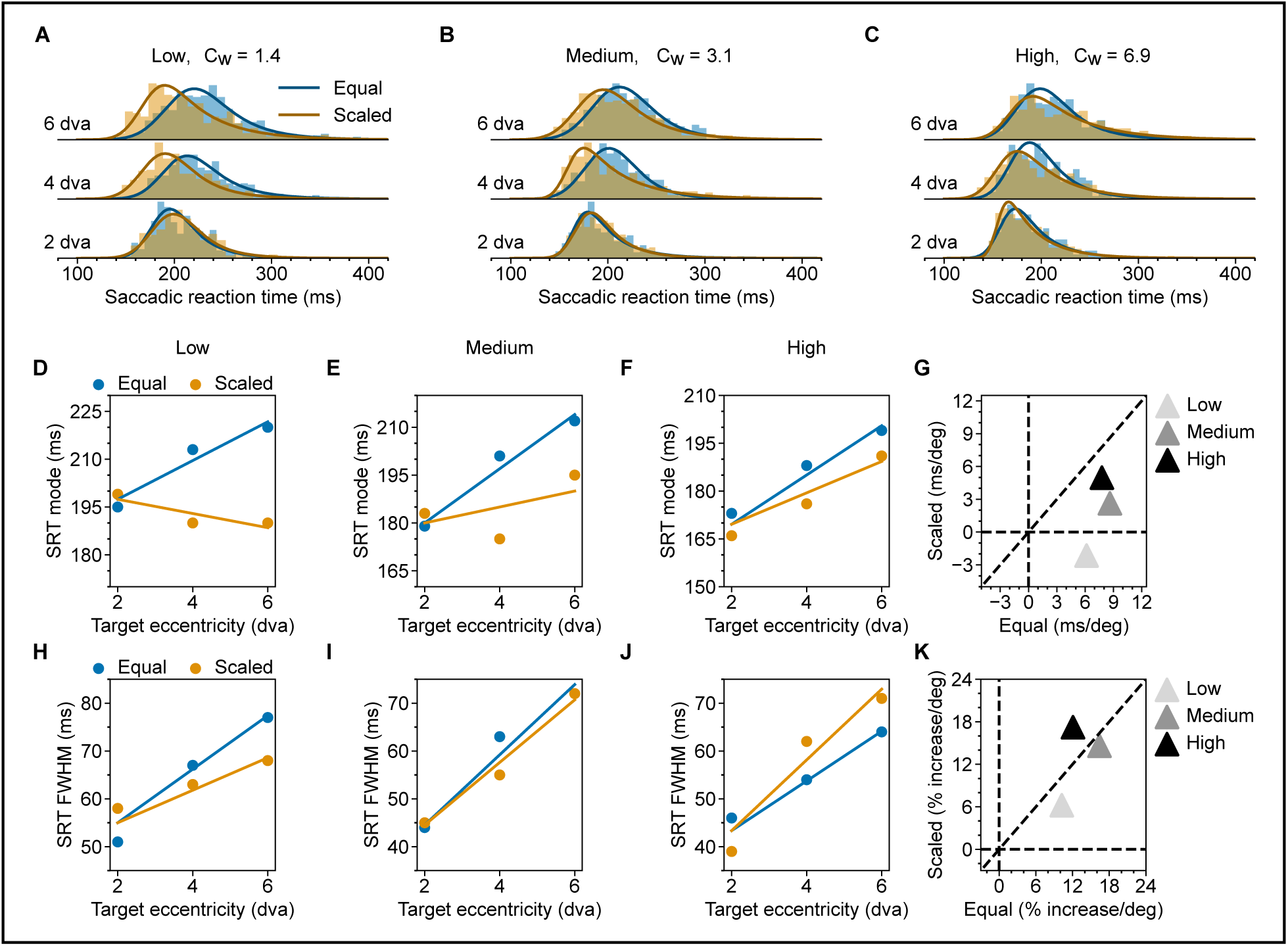
Eccentricity-dependent SRT for targets of different contrast in Monkey HO. (A-C) SRT distributions for low-, medium-, and high-contrast targets are shown separately for Equal (blue) and Scaled (orange) targets. Solid lines represent the fitted ex-Gaussian distributions. Weber contrasts, C_W_, for low-, medium-, and high-contrast targets are 1.4, 3.1, and 6.9, as also listed on top of each column. (D-F) Multiple linear regression of SRT mode on target eccentricity and scaling for the low-, medium-, and high-contrast, respectively. (G) Scatter plot of SRT mode-eccentricity slopes for Scaled vs. Equal targets at all contrast levels. (H-K) Same as (D-G), but for FWHM.

To evaluate how SRT mode depends on target eccentricity, we first fitted a simple linear regression model (*Mode* ∼ β_0_ + β_1_ ∗ (*targetEccentricity* − 2)) at each contrast level, separately for the Scaled and Equal targets. For the Equal targets, the SRT mode increased with target eccentricity at each contrast level. For the Scaled targets, the SRT mode decreased slightly with eccentricity at low contrast and increased with eccentricity at higher contrast. At each contrast level, the slopes are lower for the Scaled compared to the Equal targets. This difference in SRT-Eccentricity slopes between Scaled and Equal targets indexes the effect of scaling. To quantify the effect of scaling reducing the slope, we then fitted a multiple linear regression model (*Mode* ∼ β_0_ + β_1_ ∗ (*targetEccentricity* − 2) + β_2_ ∗ (*targetEccentricity* − 2) ∗ *IsScaled*), combining data from Equal and Scaled targets (Figure 3D-F) at each contrast level. At low and medium contrast, scaling reduced the slope significantly, but the reduction of slope due to scaling was marginal for high-contrast targets (*Low contrast*, *CI*_β2,95%_ = [−10.30, −6.00], *p* < .001; *Medium contrast*, *CI*_β2,95%_ = [−8.40, −3.80], *p* < .001; *Hig*ℎ *contrast*, *CI*_β2,95%_ = [−5.80, 1.20], *p* = .063). The above results from the multiple linear regression analysis suggest that the effect of scaling decreased at higher target contrast. This trend is illustrated graphically in Figure 3G: with increasing contrast, data points moved gradually towards the equal-slope diagonal. As before, we applied the same analysis to FWHM (Figure 3H-K). FWHM increased with more peripheral targets for both Equal and Scaled targets. Stimulus scaling had a smaller effect on FWHM. It reduced the slope only marginally for the low-contrast targets (*Low contrast*, *CI*_β2,95%_ = [−4.30, 0.50], *p* = .099; *Medium contrast*, *CI*_β2,95%_ = [−3.10, 2.30], *p* = .537; *Hig*ℎ *contrast*, *CI*_β2,95%_ = [−.60, 5.90], *p* = .171).

### Eccentricity-dependent SRT increase for endogenously driven saccade is consistent with an attentional scan from the fovea

So far, we focused on saccades in the Step task driven by the appearance of a peripheral saccade target, which occurs simultaneously with the disappearance of the fixation point; we refer to these as exogenously driven saccades. In this section, we switch to the Delayed task, where the response saccades were driven by the disappearance of the fixation point after the peripheral saccade target had already been present for 800-1500 ms, and we refer to these as endogenously driven saccades (Figure 1C). The same two stimulus sets, Equal and Scaled with filled white circles were used as target stimuli. Like the Step task, each trial in the Delayed task started with the monkey holding fixation on a fixation dot for 800 to 1000 ms. The target stimulus was then presented for 800 to 1500 ms. Importantly, during this period, the monkey was required to keep fixation on the fixation dot and restrain from making a saccade. Next, the fixation dot was turned off, which signaled the monkey to make a response saccade to the target stimulus. The monkey needed to shift its gaze to the target stimulus within 500 ms and hold its gaze on the target for another 800 to 1500 ms to complete the trial and receive a reward. Only correct trials were included in our analysis. Additionally, in 50% of the trials we included a short ≈30 ms flash at the fovea (1.0 dva diameter filled white circle) starting ≈100 ms after the fixation dot offset. The flash had no task relevance and was intended to capture attention back to the fovea when the response saccade was planned but, in most trials, not yet executed^35^. We collected data from three monkeys (KL, CH and HO) for this task.

We first present results from trials where no flash was presented. Figure 4A shows the SRT distributions from monkey KL. These SRT distributions are clearly different from the SRT distributions obtained in the Step task, wherein saccades are more reflexive. First of all, SRT in the Delayed task is generally longer. Second, SRT in the Delayed task is generally more variable. Last but not least, the effect of scaling was much smaller, with the SRT distribution for Scaled targets and Equal targets almost entirely overlapping for targets up to 8 dva. There is actually a small effect of scaling increasing SRT. All three observations are consistent with the fact that the sustained stimulus drive after the initial transient response is small and insensitive to the target size. It is worth noting that a larger stimulus may invoke lower sustained activity in the SC neurons even though its evoked transient response is larger, which agrees with the observed effect of scaling increasing SRT in the Delayed task^34^.

**Figure 4.**
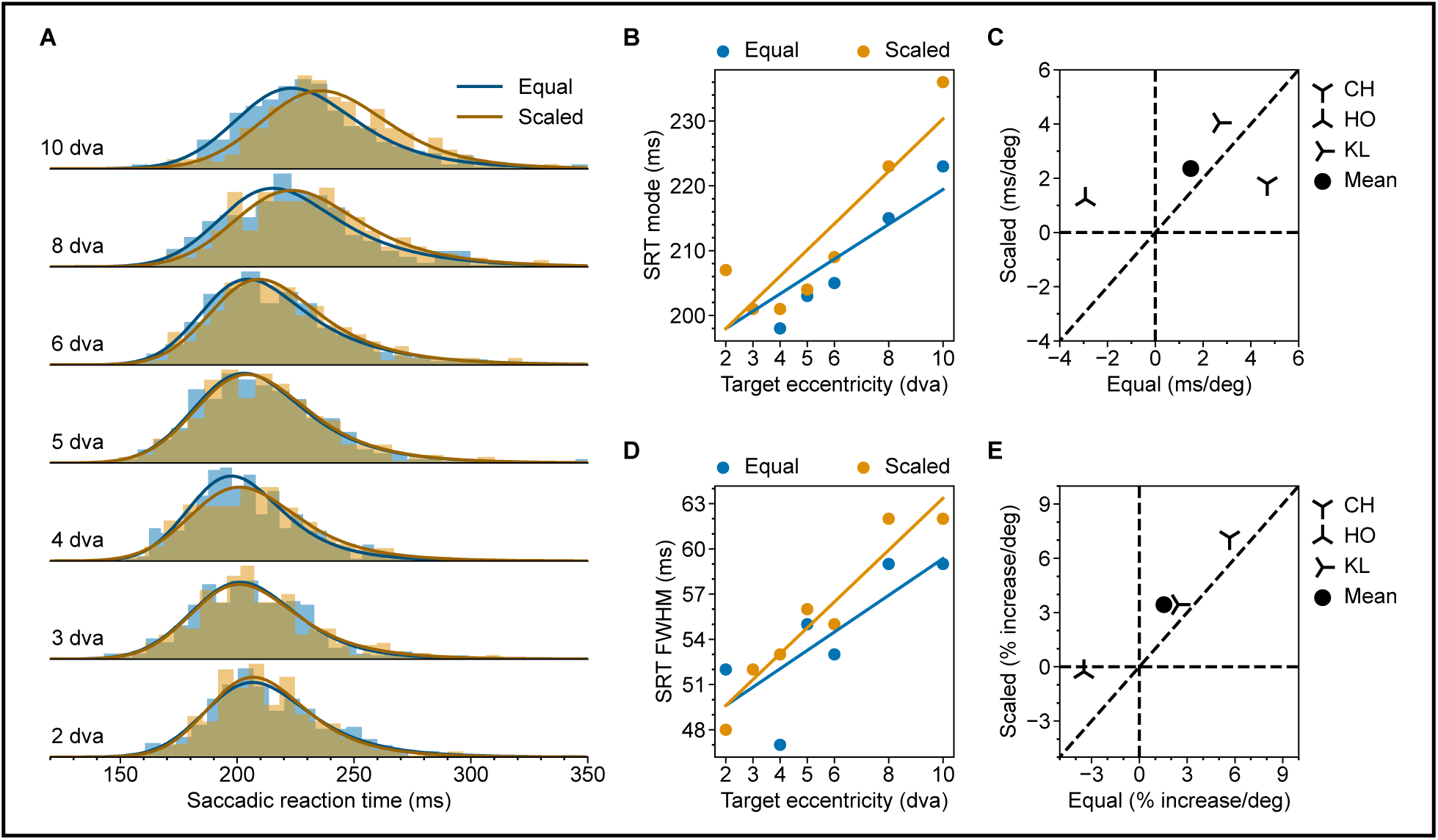
Eccentricity-dependent SRT in the Delayed task without foveal flash. (A) SRT distributions from monkey KL for Equal (Blue) and Scaled (Orange) targets. (B) Multiple linear regression of SRT mode on target eccentricity and scaling for monkey KL. (C) Scatter plot of SRT mode-eccentricity slopes for Scaled vs. Equal targets in all tested monkeys. For each monkey, the slopes are derived from the regression coefficient, as demonstrated in (B). The filled circle shows the mean slopes. (D) Same as (B) but for FWHM. (E) Same as (C) but for FWHM. The raw slopes were converted to percent increase per degree of visual angle. Results for individual monkeys other than KL are provided in the supplementary material (Figure S4-1 and S4-2).

As before, we used linear regression to quantify how the mode and FWHM of SRT distributions change with target eccentricity and stimulus scaling. Simple linear regression shows that with monkey KL, SRT mode increased with target eccentricity for both Equal and Scaled targets (*Equal*, *CI*_β1_ = [2.00, 3.22], *p* < .001; *Scaled*, *CI*_β1_ = [3.63, 4.75], *p* < .001;). Interestingly, scaling slightly increased the slope, showing the opposite effect as in the Step task (Figure 4B, *CI*_β2,95%_ = [. 85, 1.91], *p* < .001). Similar results are obtained with fixed-effect analysis pooling the three monkeys tested in this task (Figure 4C, *Equal*, 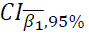 = [1.50, 2.66], *p* < .001; *Scaled*, 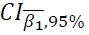 = [1.08, 2.35], *p* < .001; 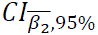 = [0.32, 1.41], *p* < .001). We repeated the same analysis for FWHM and the results is qualitatively the same as for the mode (Figure 4D, *Equal*, *CI*_β1_ = [. 51, 2.20], *p* = .009; *Scaled*, *CI*_β1_ = [. 90, 2.52], *p* < .001 Figure 4E, *Equal*, 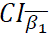 = [1.06, 2.84], *p* < .001; *Scaled*, 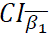= [1.99, 3.92], *p* < .001; 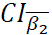= [1.03, 2.72], *p* < .001). The results for Monkey CH and HO are presented in the supplementary material (Figure S4-1 and S4-2).

The observed increase of SRT with target eccentricity in the Delayed task is intriguing. Since the delay between target onset and fixation dot offset was at least 800 ms, the monkey had enough time to prepare for the impending saccade. The increase is therefore probably not related to motor planning before the “Go” signal. Rather, it is likely related to processes that happened after the “Go” signal. Additionally, because the effect is eccentricity dependent, it is likely caused by processes that are spatially relevant. One potential candidate is attentional orienting. Note that 1) because the “Go” signal in the Delayed task is the fixation dot offset, to make a timely response, the animal needed to pay attention to the fovea, even though there were two stimuli (fixation dot and peripheral target) presented at the same time, 2) before actual saccade execution, attention is mandatorily shifted to the location of the saccade target^36^. Thus, the observed increase of SRT with target eccentricity can be explained if attentional orienting from the fovea takes longer for more eccentric targets^37^.

To provide further evidence that the eccentricity-dependent SRT increase in the Delayed task is related to attentional orienting from the fovea to the saccade target after the “GO” signal, in 50% of the trials, we presented a flash at the fovea, at ≈100 ms after the fixation-point offset (Figure 1C). When presented, this flash would drag attention back to the fovea and force a restart of the relevant processes. If the eccentricity-dependent SRT increase is indeed due to attentional orienting from the fovea to the saccade target, the additional flash stimulus would maintain this increase, albeit with a constant delay in SRT at all target locations.

Figure 5A shows the SRT distributions in trials with foveal flash from the same monkey (KL) presented in Figure 4. To facilitate comparison, we plotted the ex-Gaussian fits obtained in corresponding conditions without flashes as dashed lines. As expected, the addition of the flash prolonged the SRT^38,39^. Importantly, the eccentricity-related SRT increase persisted with the addition of the flash.

**Figure 5.**
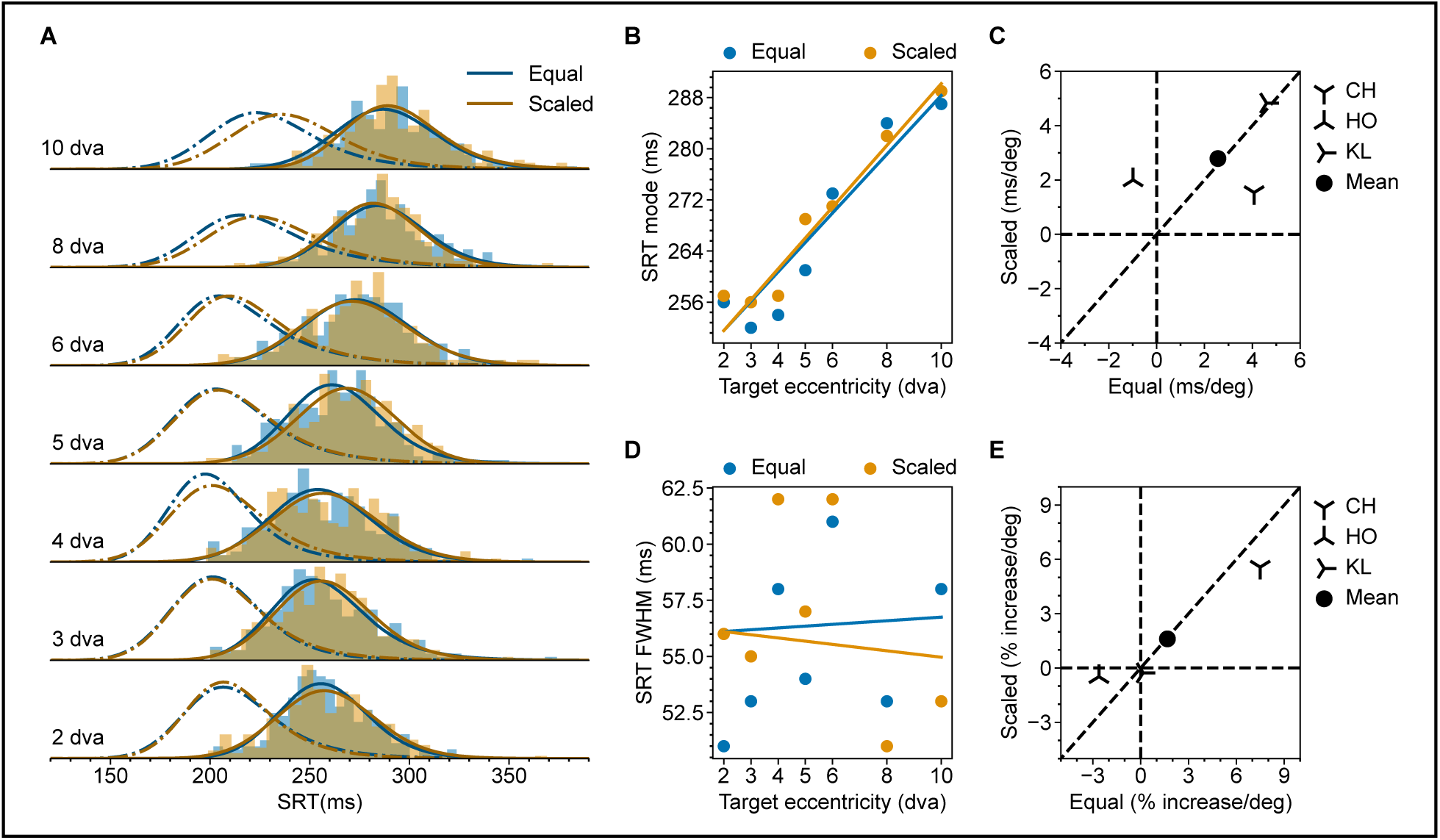
Eccentricity-dependent SRT in the Delayed task with foveal flash. (A) SRT distributions from monkey KL for Equal (Blue) and Scaled (Orange) targets. Dashed lines represent the ex-Gaussian fits for SRT distributions in corresponding conditions without the foveal flash (as in Figure 4). (B) Multiple linear regression of SRT mode on target eccentricity and scaling for monkey KL. (C) Scatter plot of SRT mode-eccentricity slopes for Scaled vs. Equal targets in all tested monkeys. For each monkey, the slopes are derived from the regression coefficient, as demonstrated in (B). The filled circle shows the mean slopes. (D) Same as (B) but for FWHM. (E) Same as (C) but for FWHM. The raw slopes were converted to percent increase per degree of visual angle. Results for individual monkeys other than KL are presented in the supplementary material (Figure S5-1 and S5-2).

We quantified these observations with linear regression. For monkey KL, SRT mode increased with target eccentricity for both Equal and Scaled targets (*Equal*, *CI*_β1_ = [4.34, 5.49], *p* < .001; *Scaled*, *CI*_β1_ = [3.98, 5.10], *p* < .001). Notably, after introducing the resetting foveal flash, which presumably further eliminated the differential activity build-up during the delay period, there is now no scaling-dependent change in slope (Figure 5B, *CI*_β2_ = [−.22, .71], *p* = .356). Intriguingly, the foveal flash also eliminated the eccentricity-dependent FWHM increase (Figure 5D, *Equal*, *CI*_β1_ = [−.32, 1.45], *p* = .223; *Scaled*, *CI*_β1_ = [−1.40, .26], *p* = .163; *CI*_β2_ = [−.91, .60], *p* = .586). The results for the other monkeys are presented in the supplementary material (Figure S5-1 and S5-2). Fixed effect analysis combining the three monkeys shows a similar pattern as described above for Monkey KL (Figure 5C, the mode: *Equal*, 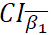 = [ 2.63, 3.50], *p* < .001; *Scaled*, 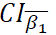= [1.88, 2.87], *p* = .016; 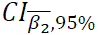 = [−.1.8, .58], *p* = .274; Figure 5E, FWHM *Equal*, 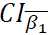= [1.14, 3.32], *p* < .001; *Scaled*, 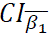 = [. 16, 2.15], *p* = .016; 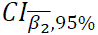= [−.90, .81], *p* = .876).

## Discussion

Saccadic eye movements play a fundamental role in organizing our spatial and temporal visual input, to the extent that SRT has become a crucial psychophysical measurement enabling us to characterize the underlying cognitive processes that would otherwise remain unseen. However, despite the importance of this measurement, some fundamental questions regarding the determinants of SRT remain, even in relatively simple situations. In this study, we examined the dependence of SRT on target eccentricity in macaque monkeys.

When designing the experiment, we based our hypotheses on previous insights into the role of the SC in generating the response saccades: Once the SC decision unit passes the response threshold, there is a relatively fixed delay in the downstream motor nuclei before the actual eye movement occurs^40,41^. Consequently, any observed dependence of SRT on target eccentricity can be attributed to the spatiotemporal activation pattern of the SC and reflects the properties of the neural processes associated with that particular task. It is these properties that have clear behavioral correlates, namely SRT, the primary interest of the current study.

The SRT reflects the total time required for target detection, response initiation and execution. A dissociation of these components, respectively their dependence on eccentricity, might have been achieved by, e.g., including a condition with a manual response that remains identical across eccentricities. Such a dissociation is beyond the scope of the present study but is an interesting target for future research.

### SRT in the Step task

In the Step task, afferent input to the SC was presumably dominated by the transient onset of the target stimulus. We first confirmed previous findings that, for medium-sized saccades, SRT increased with target eccentricity. Notably, this SRT increase was primarily explained by the foveal magnification factor in the SC (Figure 2, 3). The observed effect of foveal magnification on SRT can be explained by the increase of the accompanied RF size from the foveal to the peripheral representation in the SC. This increasing RF size compensates for the foveal magnification, thus equalizing the size of the active population for a point stimulus presented at different eccentricities, achieving a constant point image in the SC^16,42^. However, larger RFs in the periphery also mean that an equal-sized stimulus occupies a smaller proportion of the corresponding SC neurons’ RFs. The smaller occupied RF proportions likely drive the active population less strongly, leading to longer SRT^34^. When we scaled the stimulus size according to the SC magnification factor, the above-mentioned mechanism would predict an equal SRT at all eccentricities, which is what we found for the Scaled targets. This process is illustrated in Figure 6. A similar effect of foveal magnification neutralizing the eccentricity effect has been reported for a covert visual search task^43^.

**Figure 6.**
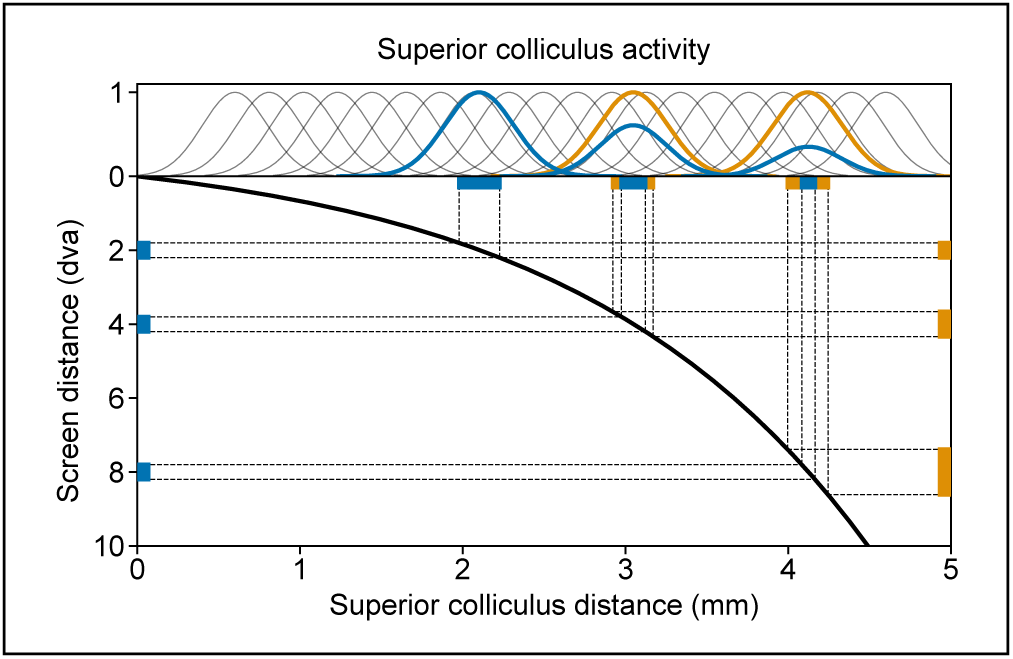
Proposed mechanism of how stimulus scaling reduces SRT in the Step task. The foveal magnification is illustrated by the thick black curve mapping points on the screen (Y-axis) to points on the SC (X-axis). Blue (orange) bars on the left (right) represent the Equal (Scaled) targets at 2, 4, and 8 degrees, respectively. The corresponding image of these stimuli in the SC is represented by horizontal blue (orange) bars at the top. These stimulus images represent the foveally-magnified afferent input to the SC. The thin gray Gaussian curves on the top illustrate example visual connectivity functions of SC neurons integrating the afferent inputs. Notice that homogenous connectivity functions, as we assumed here, correspond to RFs of increasing size with higher eccentricity when projected to the visual space. The activity level of a given SC neuron is modeled as the dot product between its Gaussian connectivity function and the afferent input. The activity profile across the SC-neuron population is calculated as the convolution between the connectivity function and the afferent signal. The thick colored curves on the top illustrate the resulting activity profiles evoked by each stimulus. Note that equally-sized stimuli (blue) activate the SC more weakly in the periphery, whereas scaled stimuli (orange) activate the SC equally strongly at all positions. Note also that the sizes of the activated neuronal populations are approximately equal because the stimuli are small compared to the SC neurons’ RFs.

Furthermore, we showed that the foveal magnification factor interacted with stimulus contrast in determining the SRT. This interaction also aligns with the known properties of neurons in the SC. Specifically, the observation that, in monkey HO, where we also modulated the target contrast, stimulus scaling modulated SRT most effectively for low-contrast stimuli can be explained by the visual response properties of SC neurons. Early recordings measuring the size tuning of SC visual responses found that with stimuli of luminance contrast 1-3 log unit higher than the response threshold, presumably corresponding to the high contrast condition in the current study, many neurons’ response magnitudes are insensitive to stimulus size. Recently, Chen and Hafed^34^ provided direct evidence for such interaction between stimulus contrast and size in driving the SC visual response. They showed that for a given SC neuron, a large stimulus evokes a larger response only if the contrast is low. When the contrast is high, the response magnitude is similar for both small and large stimuli. This interaction pattern can explain why scaling was most effective for low-contrast targets in eliminating the eccentricity-dependent SRT increase in our data (Figure 3). Together, these findings imply a direct translation of saccade target visibility (combining the effect of contrast and size) to response urgency mediated by the SC^44^.

Our analysis has been focused on comparing SRT across different eccentricities to test whether a particular kind of scaling, i.e., scaling according to the foveal magnification in the SC, affected this SRT-eccentricity relationship. Conversely, one could compare SRT at fixed target eccentricity across different scaling factors. If our hypothesis is true, that the relative afferent input strength determines the SRT-eccentricity relationship in the Step task, and that the relative size of the saccade target to the corresponding neurons’ RF can approximate this afferent input strength, one would expect that increasing the size of the saccade target at a fixed eccentricity shortens the SRT. Indeed, this expected effect of varying saccade target size on SRT has been reported before for targets that are small relative to the SC neuron’s RF size^45^. Different processes appear to be involved in saccading to very large targets^46^.

Scaling the target has an incidental consequence: a scaled target is of a different size than the fixation dot and all the non-scaled saccade targets, making it a relatively novel stimulus. Stimulus novelty can decrease reaction time^47^. However, for the difference in stimulus novelty to cause the observed effect of target scaling, i.e., flattening the SRT-eccentricity function, an additional hypothesis would be required to assign quantitatively appropriate levels of novelty to target stimuli of different sizes. Nevertheless, future research would benefit from using neutral fixation points of different types than the saccade targets to alleviate this concern^48^.

### SRT in the Delayed task

In the Delayed task, when the monkey was about to make the response saccade, the target had already been on the screen for at least 800 ms, long beyond the target-onset-related visual burst in the SC neurons, which diminishes around 100 ms after the stimulus onset^49,50^. As a result, the afferent input relevant to the response saccade was no longer expected to be dominated by the transient onset of the target stimulus. In agreement with this prediction, in the Delayed task, the dependence of SRT on eccentricity was minimally affected by scaling. Yet, the simple dependence of SRT on eccentricity was still present, leading to the question: What is the mechanism underlying this SRT increase? One potential candidate is pre-saccadic attentional shifting. Results from trials with foveal flashes provided further evidence supporting this hypothesis, as explained in the Result. If the observed SRT increase is indeed related to pre-saccadic attentional shifting, the regression slopes will inform us about how fast attention was moving from the fovea to the peripheral targets: ≈2.6 ms/dva as the average from three monkeys (Figure 5). This value, corresponding to ≈17.7 cm/s in the SC, is in a similar range as the speed of proposed attentional scanning estimated in a recent opinion paper^37^. The reported estimate was obtained partly based on neurophysiological observations and partly on a covert attention task containing a periphery reset signal. Covert attention and pre-saccadic attention are dissociable processes that can exhibit different temporal dynamics (for a recent reviewer, see Li et al.^51^). For a meaningful comparison between our results and the reported estimate, it is necessary to point out that a salient transient signal, e.g., the foveal or peripheral flash, captures attention to the spatial location of that transient in both processes^52,53^, and the moment-by-moment spatial location of attention in both processes can be determined by probing behavioral performance. These considerations help to align the observed attentional shifting speed with the estimate in Fries^37^, despite the inherent differences between covert and pre-saccadic attention.

Alternatively, the SRT increase in the Delayed task might reflect a speed-accuracy tradeoff. One way to describe the speed-accuracy tradeoff is to use the classical Fitts’s Law^54^. According to Fitts’s Law 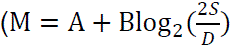 the logarithm of the ratio between movement distance (S) and target diameter (D). If we consider M to be the SRT, S to be the target eccentricity, and D to be the target size, Fitts’s Law, however, cannot explain the SRT values observed across conditions. Specifically, the fact that at the same target eccentricity, Scaled targets yielded similar or even longer SRT compared to the Equal targets is not consistent with Fitss’s Law. One might instead take D to be the size of the invisible target window, which was the same in all conditions. But then, it is difficult to explain the effect of the scaling observed in the Step task. In both cases, Fitts’s Law struggles to explain the variability of SRT in our data. Note that Wu et al.^55^ found that Fitts’s Law describes the speed-accuracy tradeoffs in a saccade sequence only when the time for the secondary, corrective saccades is included. When only the saccade latency for the primary saccade is considered as what we did in our analysis, Fitts’s law no longer holds. Furthermore, within the context of a diffusion model, the speed-accuracy tradeoff is typically modeled by adjusting the decision threshold^56^. With other parameters staying unchanged, a higher threshold implies both a longer response time and a larger variance in the response time distribution. However, we do not find that SRT mode relates consistently to FWHM. For the respective example monkey, in the Step task, scaling eliminated the eccentricity-dependent SRT mode increase but not the FWHM increase (Figure 2); in the Delayed task, adding a foveal flash eliminated the FWHM increase but not the SRT mode increase (Figure 5). All these observations indicate that a speed-accuracy tradeoff, at least in isolation, is unlikely to explain the observed SRT variation.

If the pre-saccadic attentional shift caused the SRT increase in the Delayed task, one would expect to see a similar SRT increase for medium-sized saccades in the delayed visually guided saccades task of Hafed and Goffart^4^. Indeed, although the SRT increase in their delayed task is less prominent than in their Step task, it is clearly trending. The relatively small SRT increase in the delayed task is also apparent in the present study (comparing Figures 5-6 to Figures 2-3). In addition, it is worth noting that our SRT values are generally longer than those reported by Hafed and Goffart^4^. This discrepancy is likely due to the smaller target windows used in our study (1.0 dva vs. 2-2.5 dva in radius). The size of the fixation window may be related to the speed-accuracy tradeoff as discussed earlier. Although the process involved in the speed-accuracy tradeoff may not be the direct cause of the SRT increase, it is still worth exploring how these factors interact.

Pre-saccadic attentional shifting is not expected to significantly affect SRT in the Step task. In the Step task, attention is automatically captured by the onset of the peripheral target, a process that is presumably not dependent on eccentricity. In contrast, the Delayed task involves triggering saccades by the offset of the foveal fixation dot, requiring a voluntary movement of attention from the fixation dot to the peripheral target, which we hypothesize to be eccentricity dependent.

### Variability of SRT

As noted in previous studies, SRT exhibited significant trial-by-trail variation in both the Step and Delayed tasks. Within the framework of a rise-to-threshold model, a larger variance in SRT means either a higher threshold or a more variable rate of rise^13,23,24,57^. A consistent finding in our study is the increased variability of SRT for targets located in the periphery. This pattern held true for both Equal and Scaled targets in both the Step and Delayed tasks. Notably, in the Step task with Scaled targets, SRT variance increased with target eccentricity, despite the effective flattening of the SRT mode by scaling (Figure 2, 3). These findings suggest that factors beyond foveal magnification, and to some degree invariant to the task, contributed to the observed SRT variance increase with target eccentricity. One potential mechanism could be that neuronal response fields (both motor and visual) of larger eccentricity typically have larger size. Larger response field sizes likely lead to less specific, or in other words, noisier contributions to the ensuing saccade. When cortical magnification is controlled, this leads to higher noise in the recruited neuronal population. Interestingly, Hafed and Chen^58^ reported a weaker and more variable visual response associated with less accurate saccades in SC neurons with a larger response field. Note that these authors compared neurons representing lower versus upper visual fields, yet similar properties are conceivable comparing neurons with response fields centered at different eccentricities.

### Individual differences

We observed considerable variations in SRT distributions among the four tested monkeys. These variations likely have multiple sources. First, some variance in SRT distributions could be related to the task design. Most prominently, the maximum SRT allowed was 500 ms at all locations in all tasks. This setting is essential for assuring the same task structure across all conditions despite the difference in their typical SRTs. However, this setting is not optimal for eliminating the influence of task-irrelevant processes, e.g., procrastinations to reduce the landing errors. Indeed, the monkeys whose SRTs showed the largest variance, suggesting extra noise in their decision process, were also the ones that seemed to be outliers (HU in Figure 2C and HO in Figure 4C). Since using a long response window is required to compare SRT distributions that are expected to shift significantly, future studies can use variable reward amounts to encourage the animal to make as fast a response as possible and potentially reduce the variability among the subjects^59^. Second, variations associated with scaling might be partially explained by the individual differences in foveal magnification factors^60^. Finally, it is conceivable that some of the monkeys, e.g., monkey HU, had comparatively poorer vision. Overall, the vision was sufficient for all tested monkeys to produce the observed, consistent behavior. However, subtle deficiencies in vision might have contributed to the observed individual differences, and the individual visual acuity was not tested here.

Increasing the number of subjects is crucial for obtaining a more comprehensive understanding of whether the observed effect is present in the population. However, in the current study, the number of subjects was limited, partly because of the chosen animal model, i.e., macaques. Macaque monkeys were used due to the availability of published data on the foveal magnification factor in the SC, which was essential for the experimental design. Additionally, there is a rich history of using macaque monkeys to study the underlying neural mechanisms of oculomotor control and attention. Choosing macaque monkeys thus offers the potential to use these findings to motivate future circuit investigations. In human studies, it is generally easier to include a larger number of subjects; however, there is no direct measurement of the SC foveal magnification in humans. Nevertheless, it is not unreasonable to utilize the same magnification factor as observed in macaque monkeys. Alternatively, one can consider using the magnification factor measured for the primary visual cortex, as evidence indicates that the magnification factor of the primary visual cortex and SC are similar^17^.

## Author contribution

YZ: Conceptualization, Methodology, Software, Validation, Formal Analysis, Investigation, Data Curation, Writing – Original Draft; Writing-Reviewing & Editing; PF: Methodology, Validation, Resources, Writing – Original Draft, Writing-Reviewing & Editing, Supervision, Funding Acquisition.

## Declaration of interests

P.F. has a patent on thin-film electrodes and is a member of the Advisory Board of CorTec GmbH (Freiburg, Germany). The authors declare no further competing interests.

## Declaration of Generative AI and AI-assisted technologies in the writing process

During the preparation of this work, the authors used ChatGPT and DeepL in order to enhance the manuscript’s grammatical correctness and overall readability. After using these tools, the authors reviewed and edited the content as needed and take full responsibility for the content of the publication.

## Acknowledgements

This work was supported by DFG (FOR 1847 FR2557/2-1, FR2557/5-1-CORNET, FR2557/7-1-DualStreams to P.F.), EU (HEALTH-F2-2008-200728-BrainSynch, FP7-604102-HBP to P.F.), a European Young Investigator Award to P.F., National Institutes of Health (1U54MH091657-WU-Minn-Consortium-HCP to P.F.), the LOEWE program (NeFF to P.F.).

The authors would like to thank Ziad Hafed for helpful discussions about the study, Tim Näher for advice on data analysis, Jackson Smith for training of monkey HO, and Sabrina Wallrath, Julia Hoffmann and Marianne Hartmann for technical support with monkey training and behavioral data collection.

## Figure legends

**Figure S2-1.**
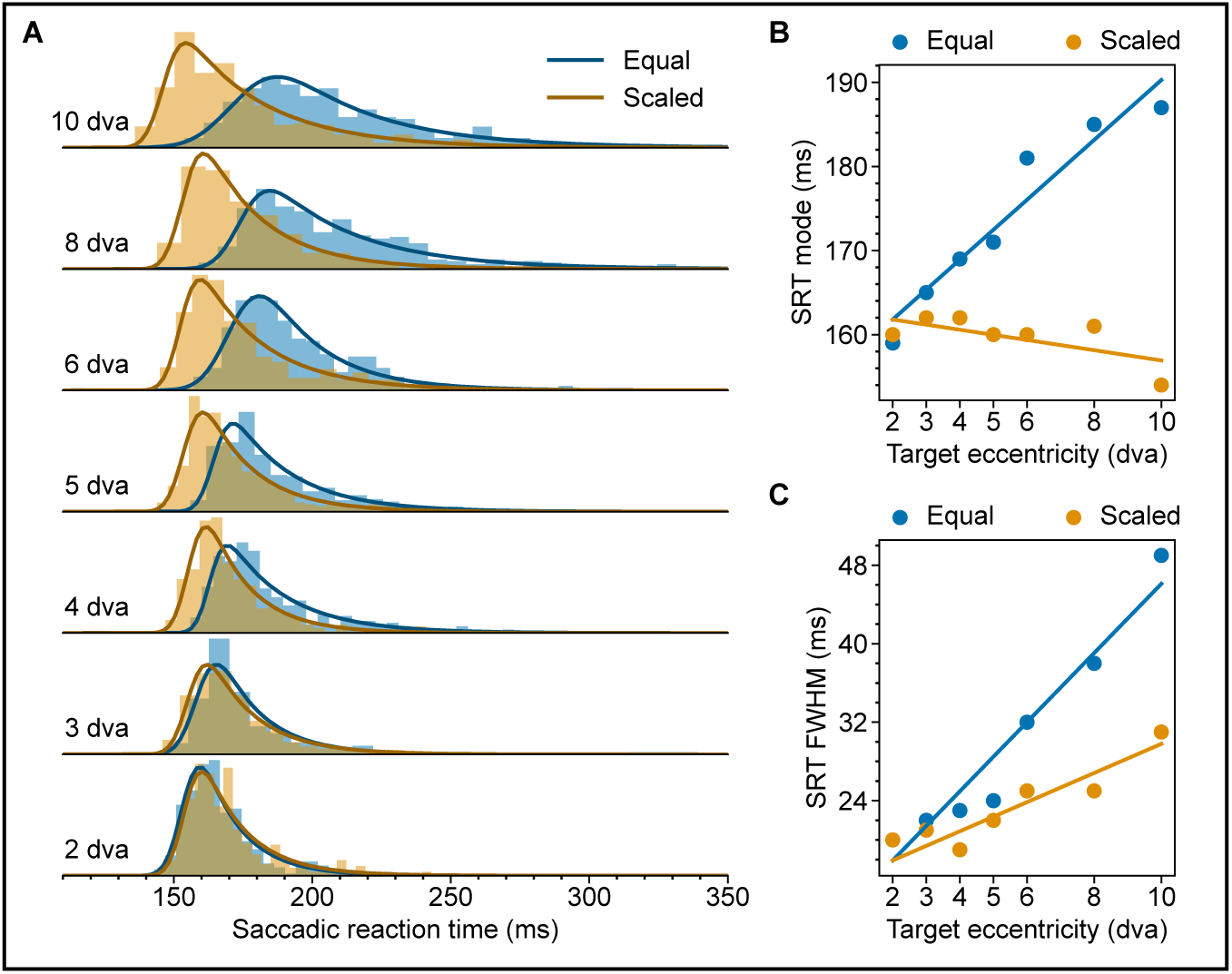
Eccentricity-dependent SRT in the Step task in Monkey CH. (A) SRT distributions for Equal (Blue) and Scaled (Orange) targets. Solid lines represent the fitted ex-Gaussian distributions. (B) Multiple linear regression of SRT mode on target eccentricity and scaling (*Equal*, *CI*_β1,95%_ = [2.99, 4.20], *p* < .001; *Scaled*, *CI*_β1,95%_ = [−0.98, −0.34], *p* < .001; *Combined*, *CI*_β2,95%_ = [−4.67, −3.72], *p* < .001). (C) Same as (B) but for FWHM (*Equal*, *CI*_β1,95%_ = [3.05, 4.46], *p* < .001; *Scaled*, *CI*_β1,95%_ = [0.91, 1.79], *p* < .001; *Combined*, *CI*_β2,95%_ = [−2.52, −1.38], *p* < .001).

**Figure S2-2.**
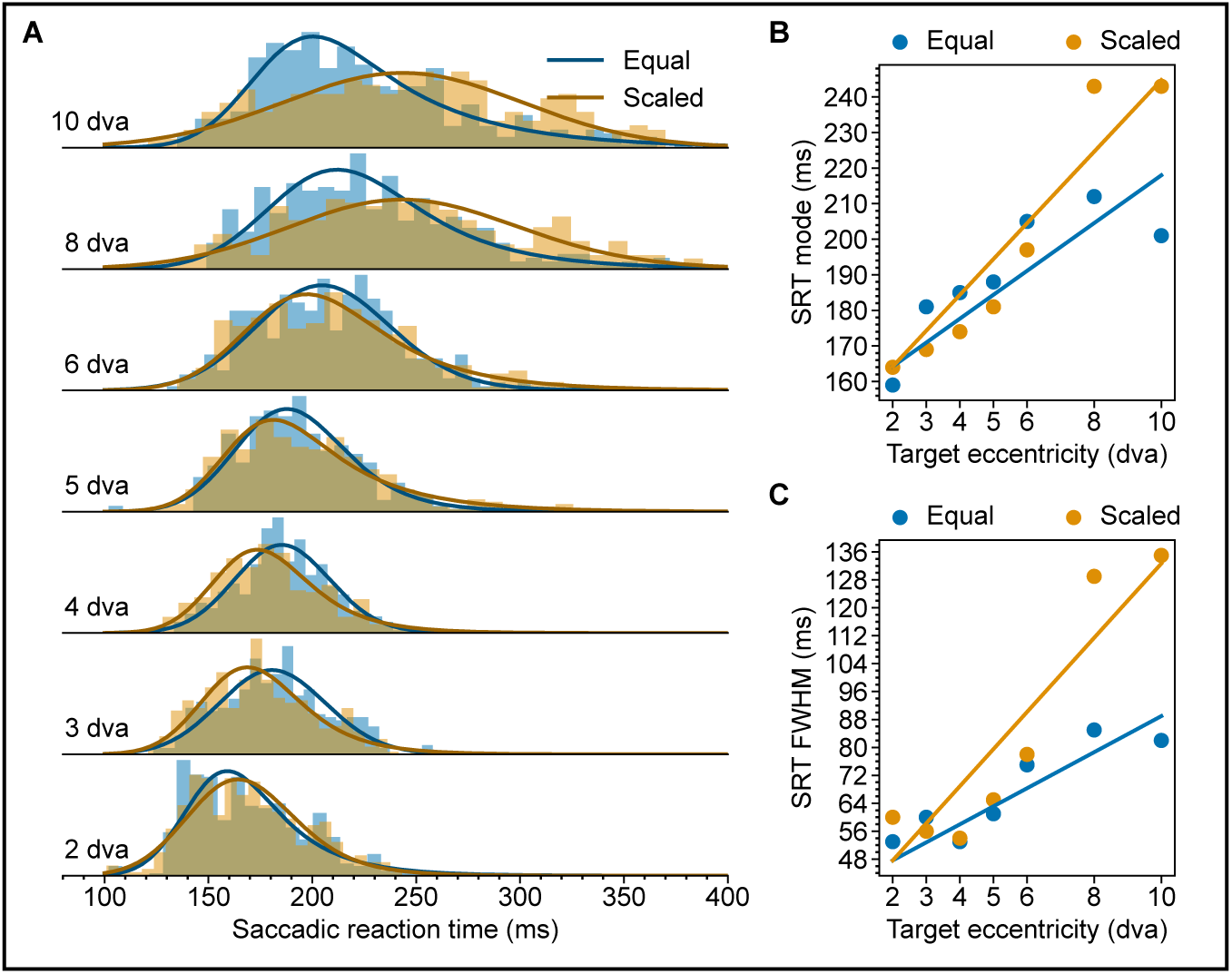
Eccentricity-dependent SRT in the Step task in Monkey HU. (A) SRT distributions for Equal (Blue) and Scaled (Orange) targets. Solid lines represent the fitted ex-Gaussian distributions. (B) Multiple linear regression of SRT mode on target eccentricity and scaling (*Equal*, *CI*_β1,95%_ = [4.06, 6.56], *p* <. .001; *Scaled*, *CI*_β1,95%_ = [10.30, 12.52], *p* < .001; *Combined*, *CI*_β2,95%_ = [2.17, 4.45], *p* < .001). (C) Same as (B) but for FWHM (*Equal*, *CI*_β1,95%_ = [3.13, 5.88], *p* < .001; *Scaled*, *CI*_β1,95%_ = [9.92, 12.62], *p* < .001; *Combined*, *CI*_β2,95%_ = [4.05, 6.55], *p* < .001).

**Figure S2-3.**
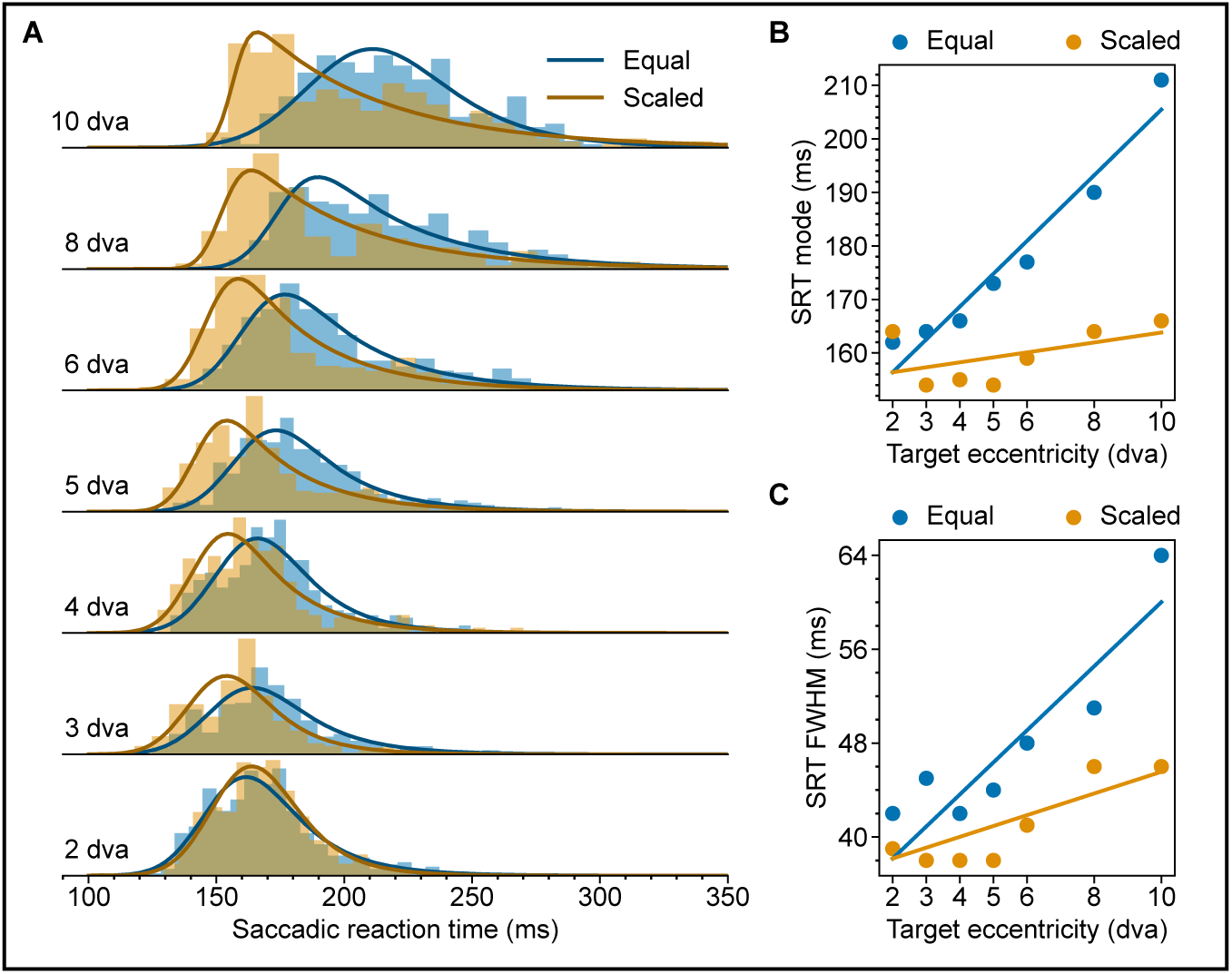
Eccentricity-dependent SRT in the Step task in Monkey KL. (A) SRT distributions for Equal (Blue) and Scaled (Orange) targets. Solid lines represent the fitted ex-Gaussian distributions. (B) Multiple linear regression of SRT mode on target eccentricity and scaling (*Equal*, *CI*_β1,95%_ = [5.04, 7.18], *p* < .001; *Scaled*, *CI*_β1,95%_ = [0.51, 1.45], *p* < .001; *Combined*, *CI*_β2,95%_ = [−6.16, −4.48], *p* <. .001). (C) Same as (B) but for FWHM (*Equal*, *CI*_β1,95%_ = [1.52, 3.69], *p* <. .001; *Scaled*, *CI*_β1,95%_ = [0.41, 1.80], *p* < .001; *Combined*, *CI*_β2,95%_ = [−2.94, −1.09], *p* = .007).

**Figure S4-1.**
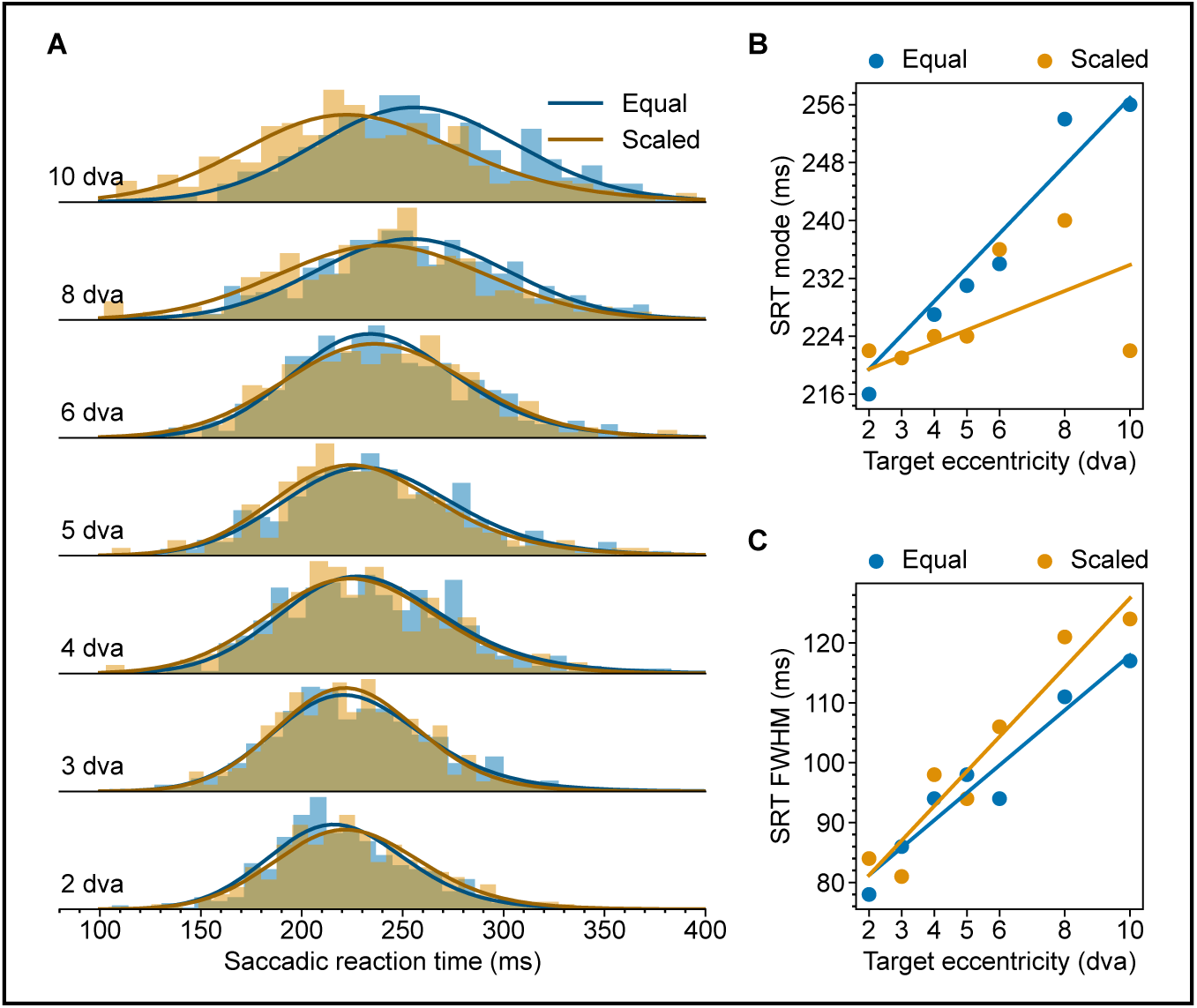
Eccentricity-dependent SRT in the Delayed task without foveal flash in Monkey CH. (A) SRT distributions for Equal (Blue) and Scaled (Orange) targets. (B) Multiple linear regression of SRT mode on target eccentricity and scaling (*Equal*, *CI*_β1,95%_ = [4.24, 6.20], *p* < .001; *Scaled*, *CI*_β1,95%_ = [. 12, 2.06], *p* = .011; *Combined*, *CI*_β2,95%_ = [−3.71, −1.83], *p* < .001). (C) Same as (B) but for FWHM (*Equal*, *CI*_β1,95%_ = [3.06, 5.67], *p* < .001; *Scaled*, *CI*_β1,95%_ = [4.30, 7.03], *p* < .001; *Combined*, *CI*_β2,95%_ = [. 06, 2.59], *p* = .053).

**Figure S4-2.**
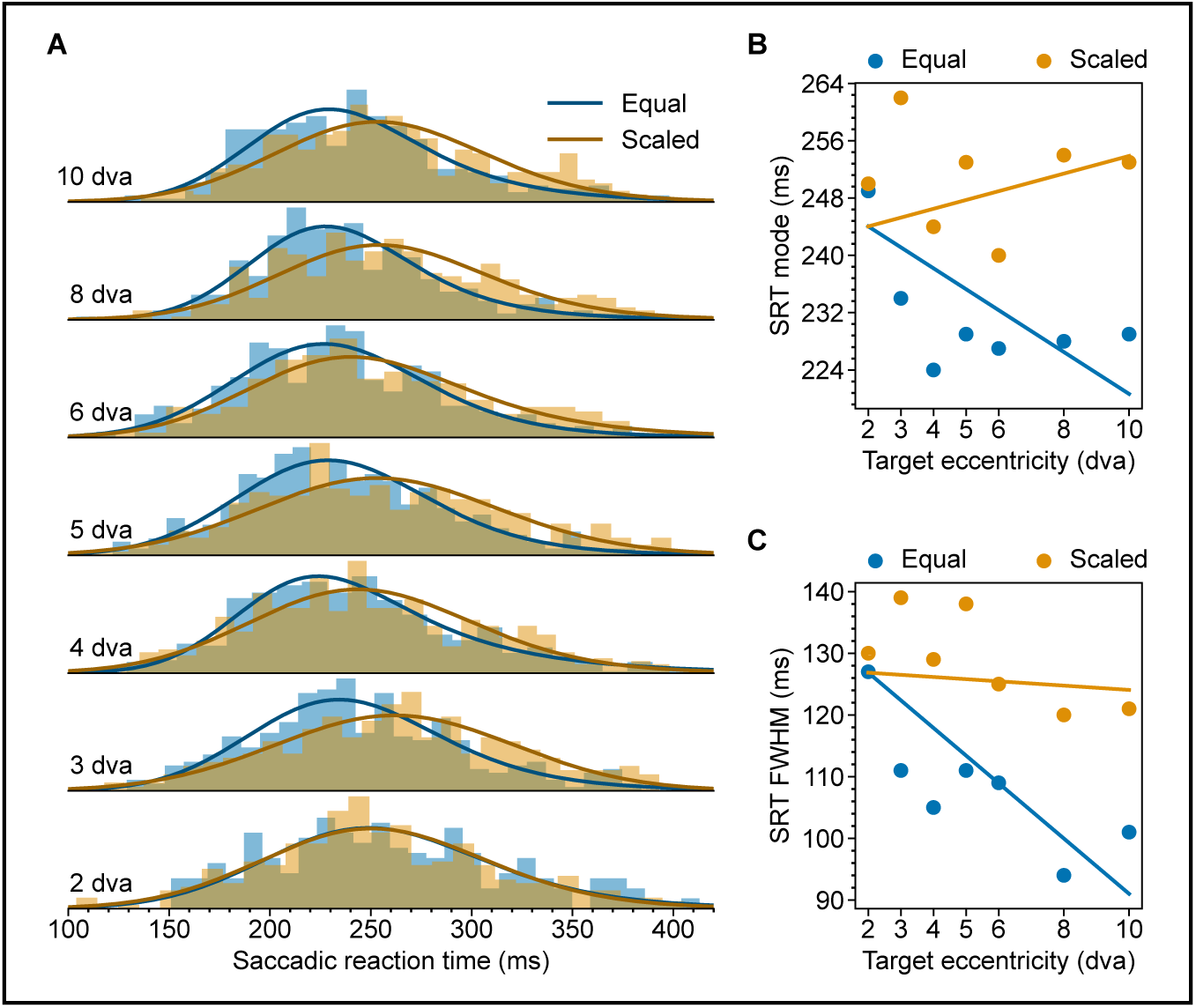
Eccentricity-dependent SRT in the Delayed task without foveal flash in Monkey HO. (A) SRT distributions for Equal (Blue) and Scaled (Orange) targets. (B) Multiple linear regression of SRT mode on target eccentricity and scaling (*Equal*, *CI*_β1,95%_ = [−3.11, −.08], *p* = .018; *Scaled*, *CI*_β1,95%_ = [−1.77, 1.41], *p* = .935; *Combined*, *CI*_β2,95%_ = [2.75, 5.19], *p* < .001). (C) Same as (B) but for FWHM (*Equal*, *CI*_β1,95%_ = [−4.73, −1.03], *p* = .001; *Scaled*, *CI*_β1,95%_ = [−4.09, −.13], *p* = .061; *Combined*, *CI*_β2,95%_ = [2.33, 5.47], *p* < .001).

**Figure S5-1.**
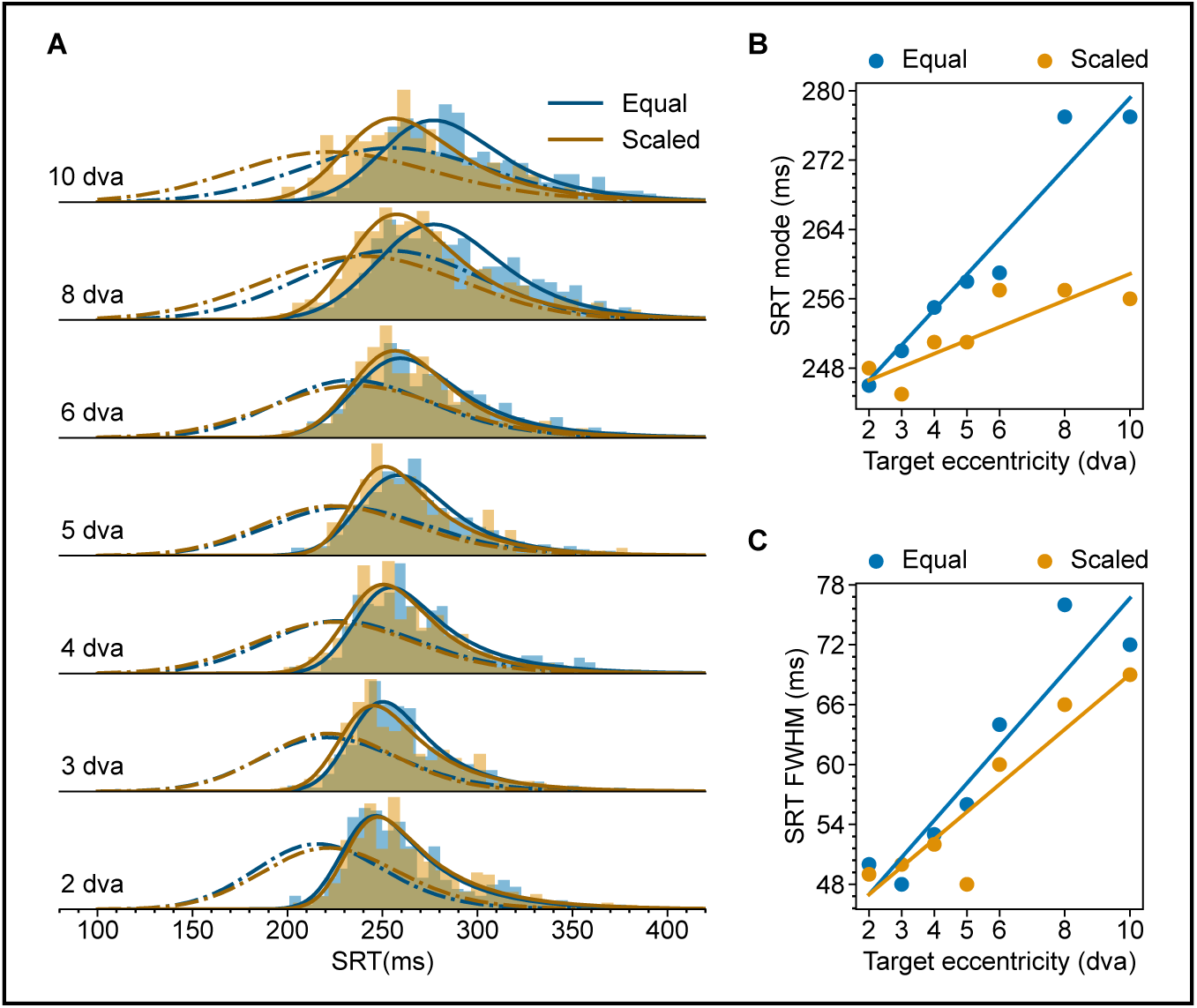
Eccentricity-dependent SRT in the Delayed task with foveal flash in Monkey CH. (A) SRT distributions for Equal (Blue) and Scaled (Orange) targets. Dashed lines represent the ex-Gaussian fits for SRT distributions in corresponding conditions without the foveal flash (as in Figure S4-1). (B) Multiple linear regression of SRT mode on target eccentricity and scaling (*Equal*, *CI*_β1,95%_ = [3.31, 5.12], *p* < .001; *Scaled*, *CI*_β1,95%_ = [. 52, 2.29], *p* < .001; *Combined*, *CI*_β2,95%_ = [−3.41, −1.81], *p* < .001). (C) Same as (B) but for FWHM (*Equal*, *CI*_β1,95%_ = [2.62, 4.85], *p* < .001; *Scaled*, *CI*_β1,95%_ = [1.82, 3.96], *p* < .001; *Combined*, *CI*_β2,95%_ = [−1.96, .12], *p* = .075).

**Figure S5-2.**
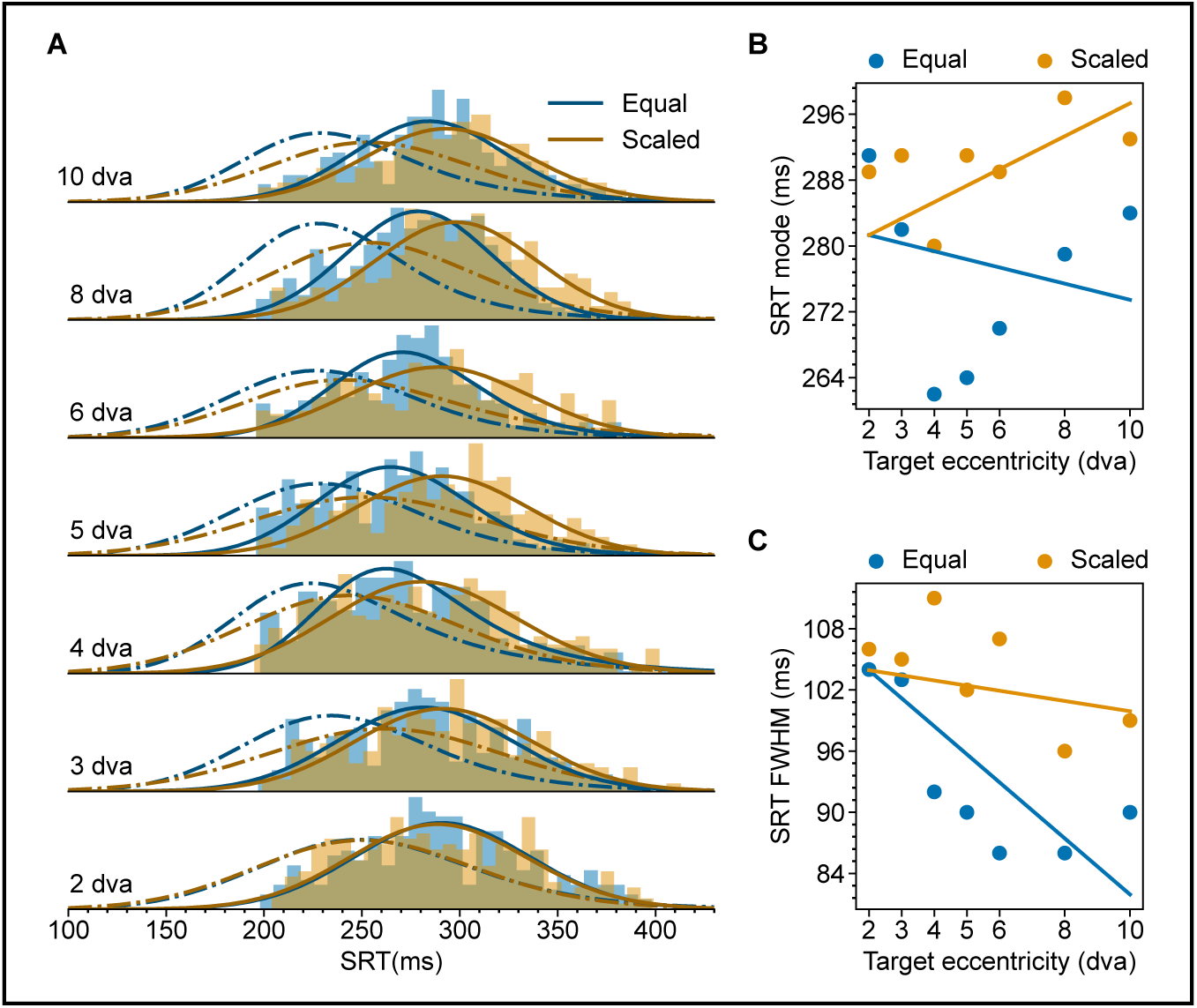
Eccentricity-dependent SRT in the Delayed task with foveal flash in Monkey HO. (A) SRT distributions for Equal (Blue) and Scaled (Orange) targets. Dashed lines represent the ex-Gaussian fits for SRT distributions in corresponding conditions without the foveal flash (as in Figure S4-2). (B) Multiple linear regression of SRT mode on target eccentricity and scaling (*Equal*, *CI*_β1,95%_ = [−.68, .93], *p* = .999; *Scaled*, *CI*_β1,95%_ = [. 201, 2.26], *p* = .012; *Combined*, *CI*_β2,95%_ = [2.38, 3.65], *p* < .001). (C) Same as (B) but for FWHM (*Equal*, *CI*_β1,95%_ = [−2.97, −0.69], *p* = .001; *Scaled*, *CI*_β1,95%_ = [−2.29, .05], *p* = .021; *Combined*, *CI*_β2,95%_ = [1.35, 3.00], *p* < .001).

## STAR Methods

### Subjects

Four male adult monkeys (Macaca mulatta) participated in the current study (referred to as CH, HO, KL and HU). All experimental procedures were conducted in compliance with the German and European animal protection laws. The experiments were approved by the responsible local authority, the Regierungspräsidium Darmstadt. All animals were implanted with a titanium headpost^61^. Additionally, Monkey KL and HO had recording chambers implanted for addressing other scientific questions. The animals’ water intake was regulated to ensure their motivation during the behavioral task in which a small juice reward was provided after each correct trial.

### Visual stimuli

All stimuli were controlled by custom software (https://github.com/esi-neuroscience/ARCADE), and presented on an LCD monitor (LG 32GK850G-B) at 143.9Hz. Viewing distance was 78 cm for all monkeys. Precise stimulus presentation time was validated with a photodiode attached to the screen. Parameters for all stimuli used are listed below.

*Fixation dot*. Trials in all tasks started with presenting a fixation dot. The fixation dot in all trials was a white filled cycle of 0.1 dva diameter, 243.7cd/m^2^, at the center of the screen.

*Target stimuli.* Each trial contained a single target stimulus chosen randomly from a predetermined set. As explained in the introduction, we focused on medium-sized saccades between 2 and 10 dva. In addition, to increase the trial counts per condition, we limited the saccade targets to the right visual field. Specifically, the stimulus set in the Step and the Delayed task contained targets centered at 2, 3, 4, 5, 6, 8, and 10 dva along the horizontal median to the right of the screen. At each target location, the set contained one target belonging to the *Equal* group and one target belonging to the *Scaled* group. For the *Equal* group, the target was a white filled circle of 243.7 cd/ m^2^ and 0.1 dva diameter at all eccentricities. For the *Scaled* group, the target was a white filled circle of 243.7 cd/ m^2^ and 0.1, 0.13, 0.17, 0.20, 0.24, 0.31, 0.38 dva diameters at 2, 3, 4, 5, 6, 8, and 10 dva, respectively. All target stimuli are small relative to the spacing between them. In the modified Step task with variable target contrast, the target locations were limited to 2, 4, and 6 dva, and at each eccentricity and for both *Scaled* and *Equal* targets, the possible target set contained targets of three different luminance levels: namely low contrast, 74.57 cd/ m^2^; medium contrast, 124.66 cd/ m^2^ and high contrast 243.7 cd/ m^2^.

*Screen background.* Gray background was used in all tasks. Except for the modified Step task with variable target contrast, the screen background luminance was 60.0 cd/ m^2^. In the modified Step task, the screen background luminance was 30.7 cd/ m^2^.

*Foveal flash.* The foveal flash used in the Delayed task was a white filled circle of 1.0 dva and 243.7 cd/ m^2^.

### Behavioral tasks

We recorded the binocular gaze data at 500 Hz with an Eyelink-1000 system (SR Research Ltd.). Real-time gaze data from one eye was used for online behavioral control. The real-time monitoring of response saccade initiation used a virtual fixation window (r ≈ 0.8 dva) centered on the fixation dot. Saccade landing on the target was determined by gaze entering and staying in a virtual target window (r ≈ 1.0 dva) centered on the saccade target. The target window radius was the same in all conditions. Additionally, the saccade flight time between leaving the fixation window and entering the target window was monitored and limited to less than ≈100 ms. In all tasks, trials of different conditions were randomly interleaved.

In the Step task (Figure 1B), after fixation was acquired and maintained for 800 to 1500 ms, the fixation dot was removed, and in the same video frame, a single target was presented in the periphery. The animal needed to make a saccade, within 500 ms, to the target and hold its gaze on the target for another 800 to 1500 ms to obtain the juice reward.

In the Delayed task (Figure 1C), after the fixation was acquired and maintained for 800 to 1000 ms, without removal of the fixation point, a single target was presented in the periphery for 800 to 1500 ms after which the fixation dot was removed. Only after fixation dot removal, the monkey was allowed to make a saccade to the target, and it needed to make a saccade towards the target within 500 ms and hold its gaze on the target for another 800 to 1500 ms to successfully complete the trial and obtain the reward. Additionally, in 50% of the trials in the Delayed task, we included a ≈30 ms flash at the fovea (white-filled circle of 1.0 dva diameter) presented ≈100 ms after the fixation dot offset. The flash had no task relevance, and the monkey needed to complete the trial as in trials without flash.

### Trial inclusion

Only correct trials were included. As described above, in these trials, the monkey made timely (SRT<500 ms) and relatively precise (error below ≈1.0 dva) primary saccades to the target. Additionally, 1) Trials with SRTs less than 100 ms were excluded from all tasks. 2) In Delayed task trials containing a foveal flash, any trial with the response saccade occurring within 100 ms after the flash was also excluded.

### Saccade detection and SRT definition

Saccades were detected using a two-dimensional velocity threshold, following the methodology described in Engbert and Kliegl^62^. In our analysis, we set lambda to 10. Saccade onset time was determined as the moment when the velocity threshold was crossed. In all tasks, SRT was defined as the time elapsed between the offset of the fixation dot and the onset of the response saccade.

### Regression analysis

The regression analysis was performed with the Python module statsmodels^63^. Further details can be found in the corresponding Results section. We used a permutation test, permuting the respective regressor label (N=1000), to test against the null hypothesis that the coefficient equals zero. The two-tailed p-values calculated from the resulting permutation distribution are reported in the main text. Bootstrapping (N=1000) was used to calculate the confidence interval. Each bootstrap sample was obtained by resampling trials within each group of trials corresponding to individual data points in the respective regression analysis.

## Notes

### Competing Interest Statement

The authors have declared no competing interest.

### Summary of Updates

Figures 1 to 5 have been revised to improve data presentation and reflect improved statistical analyses. A new Figure 6 has been added to illustrate the proposed mechanism by which stimulus scaling reduces SRT in the Step task. The discussion section has been expanded. The author details have been updated.

## References

1. Kalesnykas, R.P., and Hallett, P.E. (1994). Retinal eccentricity and the latency of eye saccades. Vision research 34, 517–531. 10.1016/0042-6989(94)90165-1.

2. Kalesnykas, R.P., and Hallett, P.E. (1996). Fixation conditions, the foveola and saccadic latency. Vision research 36, 3195–3203. 10.1016/0042-6989(96)00029-6.

3. Wyman, D., and Steinman, R.M. (1973). Letter: Latency characteristics of small saccades. Vision research 13, 2173–2175. 10.1016/0042-6989(73)90195-8.

4. Hafed, Z.M., and Goffart, L. (2020). Gaze direction as equilibrium: more evidence from spatial and temporal aspects of small-saccade triggering in the rhesus macaque monkey. Journal of neurophysiology 123, 308–322. 10.1152/jn.00588.2019.

5. Bartz, A.E. (1962). Eye-movement latency, duration, and response time as a function of angular displacement. Journal of experimental psychology 64, 318–324. 10.1037/h0043475.

6. Bell, A.H., Everling, S., and Munoz, D.P. (2000). Influence of stimulus eccentricity and direction on characteristics of pro– and antisaccades in non-human primates. Journal of neurophysiology 84, 2595–2604. 10.1152/jn.2000.84.5.2595.

7. Nuthmann, A., Vitu, F., Engbert, R., and Kliegl, R. (2016). No Evidence for a Saccadic Range Effect for Visually Guided and Memory-Guided Saccades in Simple Saccade-Targeting Tasks. PloS one 11, e0162449. 10.1371/journal.pone.0162449.

8. Pare, M., and Munoz, D.P. (1996). Saccadic reaction time in the monkey: advanced preparation of oculomotor programs is primarily responsible for express saccade occurrence. Journal of neurophysiology 76, 3666–3681. 10.1152/jn.1996.76.6.3666.

9. Dafoe, J.M., Armstrong, I.T., and Munoz, D.P. (2007). The influence of stimulus direction and eccentricity on pro– and anti-saccades in humans. Experimental brain research 179, 563–570. 10.1007/s00221-006-0817-8.

10. Land, M., Mennie, N., and Rusted, J. (1999). The roles of vision and eye movements in the control of activities of daily living. Perception 28, 1311–1328. 10.1068/p2935.

11. Bahill, A.T., Adler, D., and Stark, L. (1975). Most naturally occurring human saccades have magnitudes of 15 degrees or less. Invest Ophthalmol 14, 468–469.

12. Rohenkohl, G., Bosman, C.A., and Fries, P. (2018). Gamma Synchronization between V1 and V4 Improves Behavioral Performance. Neuron 100, 953–963 e953. 10.1016/j.neuron.2018.09.019.

13. Hanes, D.P., and Schall, J.D. (1996). Neural control of voluntary movement initiation. Science 274, 427–430. 10.1126/science.274.5286.427.

14. Liversedge, S.P., Gilchrist, I.D., and Everling, S. (2011). The Oxford handbook of eye movements (Oxford University Press).

15. Rijsdijk, J.P., Kroon, J.N., and van der Wildt, G.J. (1980). Contrast sensitivity as a function of position on the retina. Vision research 20, 235–241. 10.1016/0042-6989(80)90108-x.

16. Ottes, F.P., Van Gisbergen, J.A., and Eggermont, J.J. (1986). Visuomotor fields of the superior colliculus: a quantitative model. Vision research 26, 857–873. 10.1016/0042-6989(86)90144-6.

17. Chen, C.Y., Hoffmann, K.P., Distler, C., and Hafed, Z.M. (2019). The Foveal Visual Representation of the Primate Superior Colliculus. Current biology: CB 29, 2109–2119 e2107. 10.1016/j.cub.2019.05.040.

18. Azzopardi, P., and Cowey, A. (1996). The overrepresentation of the fovea and adjacent retina in the striate cortex and dorsal lateral geniculate nucleus of the macaque monkey. Neuroscience 72, 627–639. 10.1016/0306-4522(95)00589-7.

19. Findlay, J.M. (1980). The visual stimulus for saccadic eye movements in human observers. Perception 9, 7–21. 10.1068/p090007.

20. Cynader, M., and Berman, N. (1972). Receptive-field organization of monkey superior colliculus. Journal of neurophysiology 35, 187–201. 10.1152/jn.1972.35.2.187.

21. Chen, C.Y., Sonnenberg, L., Weller, S., Witschel, T., and Hafed, Z.M. (2018). Spatial frequency sensitivity in macaque midbrain. Nature communications 9, 2852. 10.1038/s41467-018-05302-5.

22. Doma, H., and Hallett, P.E. (1988). Rod-cone dependence of saccadic eye-movement latency in a foveating task. Vision research 28, 899–913. 10.1016/0042-6989(88)90099-5.

23. Carpenter, R.H., and Williams, M.L. (1995). Neural computation of log likelihood in control of saccadic eye movements. Nature 377, 59–62. 10.1038/377059a0.

24. Ratcliff, R., and McKoon, G. (2008). The diffusion decision model: theory and data for two-choice decision tasks. Neural computation 20, 873–922. 10.1162/neco.2008.12-06-420.

25. Otero-Millan, J., Troncoso, X.G., Macknik, S.L., Serrano-Pedraza, I., and Martinez-Conde, S. (2008). Saccades and microsaccades during visual fixation, exploration, and search: foundations for a common saccadic generator. Journal of vision 8, 21 21–18. 10.1167/8.14.21.

26. Van Zandt, T. (2000). How to fit a response time distribution. Psychon Bull Rev 7, 424–465. 10.3758/bf03214357.

27. Fries, P., and Maris, E. (2022). What to Do If N Is Two? Journal of cognitive neuroscience 34, 1114–1118. 10.1162/jocn_a_01857.

28. Lee, J., Williford, T., and Maunsell, J.H. (2007). Spatial attention and the latency of neuronal responses in macaque area V4. The Journal of neuroscience: the official journal of the Society for Neuroscience 27, 9632–9637. 10.1523/JNEUROSCI.2734-07.2007.

29. Gawne, T.J., Kjaer, T.W., and Richmond, B.J. (1996). Latency: another potential code for feature binding in striate cortex. Journal of neurophysiology 76, 1356–1360. 10.1152/jn.1996.76.2.1356.

30. Tanaka, T., Nishida, S., Aso, T., and Ogawa, T. (2013). Visual response of neurons in the lateral intraparietal area and saccadic reaction time during a visual detection task. The European journal of neuroscience 37, 942–956. 10.1111/ejn.12100.

31. Bell, A.H., Meredith, M.A., Van Opstal, A.J., and Munoz, D.P. (2006). Stimulus intensity modifies saccadic reaction time and visual response latency in the superior colliculus. Experimental brain research 174, 53–59. 10.1007/s00221-006-0420-z.

32. Pesaran, B., Vinck, M., Einevoll, G.T., Sirota, A., Fries, P., Siegel, M., Truccolo, W., Schroeder, C.E., and Srinivasan, R. (2018). Investigating large-scale brain dynamics using field potential recordings: analysis and interpretation. Nature neuroscience 21, 903–919. 10.1038/s41593-018-0171-8.

33. Ludwig, C.J., Gilchrist, I.D., and McSorley, E. (2004). The influence of spatial frequency and contrast on saccade latencies. Vision research 44, 2597–2604. 10.1016/j.visres.2004.05.022.

34. Chen, C.Y., and Hafed, Z.M. (2018). Orientation and Contrast Tuning Properties and Temporal Flicker Fusion Characteristics of Primate Superior Colliculus Neurons. Frontiers in neural circuits 12, 58. 10.3389/fncir.2018.00058.

35. Nakayama, K., and Mackeben, M. (1989). Sustained and transient components of focal visual attention. Vision research 29, 1631–1647. 10.1016/0042-6989(89)90144-2.

36. Deubel, H., and Schneider, W.X. (1996). Saccade target selection and object recognition: evidence for a common attentional mechanism. Vision research 36, 1827–1837. 10.1016/0042-6989(95)00294-4.

37. Fries, P. (2023). Rhythmic attentional scanning. Neuron 111, 954–970. 10.1016/j.neuron.2023.02.015.

38. Buonocore, A., and McIntosh, R.D. (2008). Saccadic inhibition underlies the remote distractor effect. Experimental brain research 191, 117–122. 10.1007/s00221-008-1558-7.

39. Reingold, E.M., and Stampe, D.M. (2002). Saccadic inhibition in voluntary and reflexive saccades. Journal of cognitive neuroscience 14, 371–388. 10.1162/089892902317361903.

40. Schall, J.D., and Paré, M. (2021). The unknown but knowable relationship between Presaccadic Accumulation of activity and Saccade initiation. Journal of computational neuroscience 49, 213–228. 10.1007/s10827-021-00784-7.

41. Gandhi, N.J., and Katnani, H.A. (2011). Motor functions of the superior colliculus. Annual review of neuroscience 34, 205–231. 10.1146/annurev-neuro-061010-113728.

42. Harvey, B.M., and Dumoulin, S.O. (2011). The relationship between cortical magnification factor and population receptive field size in human visual cortex: constancies in cortical architecture. The Journal of neuroscience: the official journal of the Society for Neuroscience 31, 13604–13612. 10.1523/JNEUROSCI.2572-11.2011.

43. Carrasco, M., and Frieder, K.S. (1997). Cortical magnification neutralizes the eccentricity effect in visual search. Vision research 37, 63–82. 10.1016/s0042-6989(96)00102-2.

44. Veale, R., Hafed, Z.M., and Yoshida, M. (2017). How is visual salience computed in the brain? Insights from behaviour, neurobiology and modelling. Philosophical transactions of the Royal Society of London. Series B, Biological sciences 372. 10.1098/rstb.2016.0113.

45. Boch, R., Fischer, B., and Ramsperger, E. (1984). Express-saccades of the monkey: reaction times versus intensity, size, duration, and eccentricity of their targets. Experimental brain research 55, 223–231. 10.1007/BF00237273.

46. De Vries, J.P., Azadi, R., and Harwood, M.R. (2016). The saccadic size-latency phenomenon explored: Proximal target size is a determining factor in the saccade latency. Vision research 129, 87–97. 10.1016/j.visres.2016.09.006.

47. Koster, R., Seow, T.X., Dolan, R.J., and Duzel, E. (2016). Stimulus Novelty Energizes Actions in the Absence of Explicit Reward. PloS one 11, e0159120. 10.1371/journal.pone.0159120.

48. Thaler, L., Schutz, A.C., Goodale, M.A., and Gegenfurtner, K.R. (2013). What is the best fixation target? The effect of target shape on stability of fixational eye movements. Vision research 76, 31–42. 10.1016/j.visres.2012.10.012.

49. Hafed, Z.M., Hoffmann, K.P., Chen, C.Y., and Bogadhi, A.R. (2023). Visual Functions of the Primate Superior Colliculus. Annual review of vision science. 10.1146/annurev-vision-111022-123817.

50. Goldberg, M.E., and Wurtz, R.H. (1972). Activity of superior colliculus in behaving monkey. I. Visual receptive fields of single neurons. Journal of neurophysiology 35, 542–559. 10.1152/jn.1972.35.4.542.

51. Li, H.H., Hanning, N.M., and Carrasco, M. (2021). To look or not to look: dissociating presaccadic and covert spatial attention. Trends in neurosciences 44, 669–686. 10.1016/j.tins.2021.05.002.

52. Theeuwes, J., Kramer, A.F., Hahn, S., Irwin, D.E., and Zelinsky, G.J. (1999). Influence of attentional capture on oculomotor control. J Exp Psychol Hum Percept Perform 25, 1595–1608. 10.1037//0096-1523.25.6.1595.

53. Remington, R.W., Johnston, J.C., and Yantis, S. (1992). Involuntary attentional capture by abrupt onsets. Percept Psychophys 51, 279–290. 10.3758/bf03212254.

54. Fitts, P.M. (1954). The information capacity of the human motor system in controlling the amplitude of movement. Journal of experimental psychology 47, 381–391.

55. Wu, C.C., Kwon, O.S., and Kowler, E. (2010). Fitts’s Law and speed/accuracy trade-offs during sequences of saccades: Implications for strategies of saccadic planning. Vision research 50, 2142–2157. 10.1016/j.visres.2010.08.008.

56. Ratcliff, R., Smith, P.L., Brown, S.D., and McKoon, G. (2016). Diffusion Decision Model: Current Issues and History. Trends in cognitive sciences 20, 260–281. 10.1016/j.tics.2016.01.007.

57. Trappenberg, T.P., Dorris, M.C., Munoz, D.P., and Klein, R.M. (2001). A model of saccade initiation based on the competitive integration of exogenous and endogenous signals in the superior colliculus. Journal of cognitive neuroscience 13, 256–271. 10.1162/089892901564306.

58. Hafed, Z.M., and Chen, C.Y. (2016). Sharper, Stronger, Faster Upper Visual Field Representation in Primate Superior Colliculus. Current biology: CB 26, 1647–1658. 10.1016/j.cub.2016.04.059.

59. Fischer, B., and Wegener, D. (2018). Emphasizing the “positive” in positive reinforcement: using nonbinary rewarding for training monkeys on cognitive tasks. Journal of neurophysiology 120, 115–128. 10.1152/jn.00572.2017.

60. Van Essen, D.C., Newsome, W.T., and Maunsell, J.H. (1984). The visual field representation in striate cortex of the macaque monkey: asymmetries, anisotropies, and individual variability. Vision research 24, 429–448. 10.1016/0042-6989(84)90041-5.

61. Psarou, E., Vezoli, J., Schölvinck, M.L., Ferracci, P.-A., Zhang, Y., Grothe, I., Roese, R., and Fries, P. (2022). Modular, cement-free, customized headpost and connector-chamber implants for macaques. bioRxiv, 2022.2011.2009.515849. 10.1101/2022.11.09.515849%JbioRxiv.

62. Engbert, R., and Kliegl, R. (2003). Microsaccades uncover the orientation of covert attention. Vision research 43, 1035–1045. 10.1016/s0042-6989(03)00084-1.

63. Seabold, Skipper, and Perktold, J. (2010). statsmodels: Econometric and statistical modeling with python.

